# Cohesin-mediated DNA loop extrusion resolves sister chromatids in G2 phase

**DOI:** 10.1101/2023.01.12.523718

**Authors:** Paul Batty, Christoph C.H. Langer, Zsuzsanna Takács, Wen Tang, Claudia Blaukopf, Jan-Michael Peters, Daniel W. Gerlich

**Affiliations:** Institute of Molecular Biotechnology of the Austrian Academy of Sciences (IMBA), Vienna BioCenter (VBC), 1030 Vienna, Austria; Vienna BioCenter PhD Program, Doctoral School of the University of Vienna and Medical University of Vienna, A-1030, Vienna, Austria; Research Institute of Molecular Pathology (IMP), Vienna BioCenter (VBC), Vienna, 1030 Vienna, Austria

## Abstract

Genetic information is stored in linear DNA molecules, which fold extensively inside cells. DNA replication along the folded template path yields two sister chromatids that initially occupy the same nuclear region in a highly intertwined arrangement. Dividing cells must disentangle and condense the sister chromatids into separate bodies such that a microtubule-based spindle can move them to opposite poles. While the spindle-mediated transport of sister chromatids has been studied in detail, the chromosome-intrinsic mechanics pre-segregating sister chromatids have remained elusive. Here, we show that human sister chromatids resolve extensively already during interphase, in a process dependent on the loop-extruding activity of cohesin, but not that of condensins. Increasing cohesin’s looping capability increases sister DNA resolution in interphase nuclei to an extent normally seen only during mitosis, despite the presence of abundant arm cohesion. That cohesin can resolve sister chromatids so extensively in the absence of mitosis-specific activities indicates that DNA loop extrusion is a generic mechanism for segregating replicated genomes, shared across different Structural Maintenance of Chromosomes (SMC) protein complexes in all kingdoms of life.

## Introduction

Chromosomes reorganise extensively during the cell cycle to enable gene expression during interphase and chromosome segregation during cell division. This dynamic reorganisation of chromosomes during the cell cycle is regulated by various SMC protein complexes (van Ruiten & Rowland, 2018; Batty & Gerlich, 2019; Yatskevich *et al*, 2019; Davidson & Peters, 2021; Mirny & Dekker, 2022) which establish topological linkages between DNA molecules (Gruber *et al*, 2003; Ivanov & Nasmyth, 2005) and extrude DNA into loops through a motor activity (Ganji *et al*, 2018; Davidson *et al*, 2019; Kim *et al*, 2019; Golfier *et al*, 2020).

The SMC protein complex cohesin was originally discovered as a factor linking sister chromatids along their arms (Michaelis *et al*, 1997; Guacci *et al*, 1997) through the topological embrace of two DNA molecules (Gruber *et al*, 2003; Ivanov & Nasmyth, 2005; Haering *et al*, 2008; Srinivasan *et al*, 2018). A distinct subset of cohesin complexes was later found to extrude DNA into dynamic loops (Davidson *et al*, 2019; Kim *et al*, 2019; Golfier *et al*, 2020) throughout interphase to fold chromosomes of vertebrate cells into topologically associating domains (TADs) (Rao *et al*, 2017; Schwarzer *et al*, 2017; Wutz *et al*, 2017; Gassler *et al*, 2017). Most cohesin dissociates from chromosome arms during mitotic entry in vertebrate cells (Waizenegger *et al*, 2000; Gerlich *et al*, 2006b; Liang *et al*, 2015) in a process that depends on the protein WAPL (Gandhi *et al*, 2006; Kueng *et al*, 2006). During mitosis the structurally related SMC protein complexes, condensin I and condensin II, associate with chromosomes (Ono *et al*, 2003; Hirota *et al*, 2004; Gerlich *et al*, 2006a; Walther *et al*, 2018) to form DNA loops in mitotic cells that are larger than those formed by cohesin in interphase cells (Naumova *et al*, 2013; Gibcus *et al*, 2018). How these cell cycle-regulated activities of cohesin and condensin contribute to sister chromatid resolution, however, has remained unclear.

Imaging chromosomes with differentially labelled sister DNAs suggests that sister chromatids begin to resolve at the onset of mitotic prophase (Nagasaka *et al*, 2016) through a process that depends on condensins, topoisomerase II, and WAPL (Gandhi et al, 2006; Kueng et al, 2006; Nagasaka et al, 2016; Eykelenboom & Gierli, 2019)These observations support a model in which removal of arm cohesion combined with an increase in DNA loop-extruding activity by condensin in the presence of a DNA strand passaging activity is necessary and sufficient to promote sister chromatid resolution (Nasmyth, 2001; Goloborodko *et al*, 2016a). Imaging fluorescently labelled genomic loci, however, showed that sister chromatids transiently split already during interphase, often as far apart as in mitotic cells (Ono *et al*, 2013; Stanyte *et al*, 2018; Eykelenboom *et al* 2019). Moreover, sister chromatid-sensitive chromosome conformation capture (scsHi-C) revealed that sister chromatids locally separate within TADs, despite abundant chromosome arm cohesion at this cell cycle stage (Mitter *et al*, 2020). Together, these findings raise the question as to how arm cohesion affects the resolution of sister chromatids, and how cohesin-mediated loop extrusion compares to condensin-mediated loop extrusion in its capacity to promote sister chromatid resolution. As previous structural analyses of sister chromatids were based on very harsh DNA denaturation and fixation procedures (Nagasaka *et al*, 2016) that are known to disrupt the structure of chromosomes (Beckwith *et al*, 2022), we aimed to develop improved methodologies to study the fine structure of sister chromatids in human cells not only in mitosis, but also during interphase.

## Results

### Cohesin resolves sister chromatids in G2

To implement an improved method for the detection of sister chromatid resolution, we used the nucleotide analogue F-ara-EdU, as it has low toxicity and can be visualised without DNA denaturation (Salic & Mitchison, 2008). To generate cells in which each replicated chromosome contains a single labelled sister chromatid, we cultured synchronised HeLa cells for one S phase in the presence of F-ara-EdU, before subsequently removing F-ara-EdU and letting the cells divide and progress further through one additional S phase (Fig. 1a, Fig. EV. 1a). We then visualised F-ara-EdU by attaching a fluorophore using click-chemistry and imaged cells by AiryScan microscopy.

**Figure 1.**
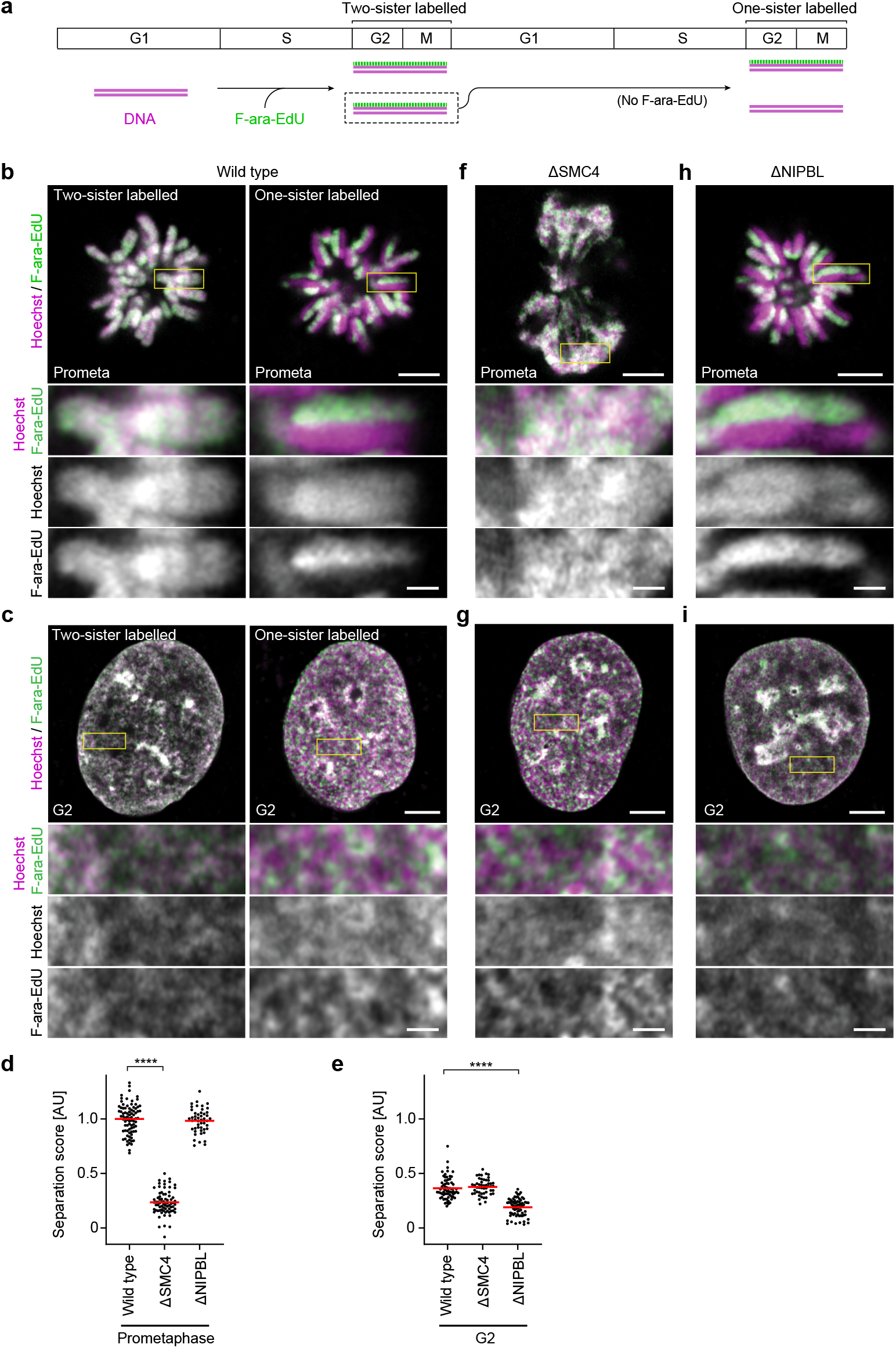
Cohesin resolves sister chromatids in G2. **a**, Sister chromatid labelling assay. Cells pre-synchronised to the G1/S boundary are cultured for one cell cycle in the presence of F-ara-EdU and then for another cell cycle in nucleotide-free medium to obtain cells with one sister chromatid labelled on each replicated chromosome. Cells with two sister chromatids labelled per chromosome are generated by fixing the cells after culturing for one cell cycle in F-ara-EdU. For analysis, cells are arrested either in prometaphase by STLC or in G2 by RO-3306 and DNA is stained using Hoechst 33342. F-ara-EdU is visualised through conjugation of AF-488. **b**, Representative images from prometaphase cells labelled on one or two sister chromatids as indicated. **c**, Representative images from G2 cells labelled on one or two sister chromatids as indicated. **d**, Quantification of sister chromatid separation in one-sister labelled prometaphase cells for experimental conditions as indicated. Dots represent individual cells; red bars indicate the mean. Wild type (n = 83 cells, from 14 experimental replicates), ΔSMC4 (n = 75 cells, from 9 experimental replicates) and ΔNIPBL (n = 48 cells, from 4 experimental replicates) were analysed. Significance was tested using a two-tailed Mann Whitney U test; P = 2.33 × 10^−27^. **e**, Quantification of sister chromatid separation as in **d** for one-sister labelled G2 cells. Wild type (n = 69 cells, from 8 experimental replicates), ΔSMC4 (n = 50 cells from 5 experimental replicates) and ΔNIPBL (n = 70 cells from 7 experimental replicates) were analysed. Significance was tested using a two-tailed Mann Whitney U test; P = 6.55 × 10^−19^. **f**, Representative images from ΔSMC4 prometaphase cells labelled on one sister chromatid. **g**, Representative images from ΔSMC4 G2 cells labelled on one sister chromatid. **h**, Representative images from ΔNIPBL prometaphase cells labelled on one sister chromatid. **i**, Representative images from ΔNIPBL G2 cells labelled on one sister chromatid. All images show single Z-slices from 3D-stacks. Yellow boxes indicate inset regions. Scale bars large panels: 5 µm, insets: 1 µm.

To validate our approach, we first assessed the efficiency and specificity of sister chromatid labelling. In the first G2 phase and mitosis after culturing cells in F-ara-EdU, we observed a strong overlap between F-ara-EdU fluorescence and the general DNA dye Hoechst 33342, such that mitotic chromosome arms were labelled on both sister chromatids (Fig. 1b, two-sister labelled; Fig. EV. 1b). Mitotic cells that had progressed through an additional S phase in the absence of F-ara-EdU had chromosome arms labelled only on a single sister (Fig. 1b, one-sister labelled; Fig. EV. 1c). Thus, our procedure specifically and completely labels one sister chromatid per chromosome.

To quantify the resolution of sister chromatids we developed an automated analysis pipeline to calculate the pixel-wise correlation between F-ara-EdU fluorescence and Hoechst 33342 (Fig. EV. 1d) (see Materials and Methods for details). In the first mitosis after labelling, where a high degree of colocalisation between Hoechst and F-ara-EdU is expected, the Spearman correlation coefficient was close to 1, demonstrating homogeneous and complete labeling of both sister chromatids. In the second mitosis, after cells had progressed through one S phase in the absence of F-ara-EdU, the Spearman correlation coefficient dropped to 0.5, indicating extensive resolution of sister chromatids (Fig. EV. 1e-g). Based on these two reference conditions, we calculated a normalised sister chromatid separation score (Fig. 1d, e), thereby allowing investigation of sister chromatid resolution under various experimental conditions.

We first assessed the organisation of sister chromatids in wild type cells synchronised to G2 by the CDK1 inhibitor RO-3306 (Vassilev *et al*, 2006). Imaging cells with single sister chromatids labelled revealed substantial separation of F-ara-EdU and Hoechst 33342 that was not visible in G2 cells with both sister chromatids labelled (Fig. 1c, e). By immunofluorescence detection of histone 3 phosphorylation and cyclin B1 localisation we confirmed that these cells had indeed not entered mitosis (Fig. EV. 2a-d). Analysis of wild type cells without RO-3306 treatment gave similar results (Fig. EV. 2e-f), showing that such separation was not due to secondary effects resulting from CDK1 inhibition. Thus, even though sister chromatids do not form discrete bodies in interphase nuclei, they resolve extensively before cells enter mitosis.

The resolution of sister chromatids in G2 might be mediated by partial activation of condensin II, which binds to chromatin already prior to mitotic entry (Hirota *et al*, 2004; Gerlich *et al*, 2006a; Ono *et al*, 2013). To test whether the resolution of sister chromatids in G2 cells depends on condensin, we used auxin-induced degradation to deplete SMC4, an essential component of the condensin complex, such that cells progressed through S and G2 phase in the absence of condensins (Fig. EV. 3a-d). Condensin depletion completely abrogated the formation of resolved cylindrical chromosomes in the following mitosis (Fig. 1d, f), yet it had no detectable effect on the resolution of sister chromatids in G2 cells (Fig. 1e, g). Thus, while our data confirm that condensin is an essential factor for sister chromatid resolution in mitosis (Nagasaka *et al*, 2016), the pre-mitotic resolution of sister chromatids in G2 nuclei must be driven by other factors.

Given that cohesin continuously forms DNA loops throughout interphase (Rao *et al*, 2017; Schwarzer *et al*, 2017; Wutz *et al*, 2017; Gassler *et al*, 2017), we investigated its role in sister chromatid resolution. To determine the effect of cohesin-mediated DNA looping we used auxin-mediated degradation to deplete NIPBL, a cofactor that is essential for cohesin-mediated loop extrusion *in vitro* (Davidson *et al*, 2019; Kim *et al*, 2019) and cohesin-mediated looping in cells (Schwarzer *et al*, 2017; Mitter *et al*, 2020). By adding the auxin analogue 5-Ph-IAA after cells had progressed through S phase, we induced NIPBL depletion to levels undetectable by Western blotting in G2-synchronised cells (Fig. EV. 3e-h). While the AID-tagged cell line showed reduced NIPBL levels already prior to 5-Ph-IAA addition compared to wild type cells, previous characterisation has shown that this residual background degradation does not impair sister chromatid cohesion (Mitter *et al*, 2020). The depletion of NIPBL substantially reduced sister chromatid resolution in G2 cells, yet in the subsequent mitosis sister chromatids were fully resolved (Fig. 1d, e, h, i). Thus, an initial resolution of sister chromatids during interphase depends on cohesin-mediated DNA looping, whereas in the subsequent mitosis condensin takes over to promote more extensive resolution into cylindrical sister chromatids.

### Hyperactive cohesin forms mitosis-like sister chromatids in G2

That sister chromatids resolve less in G2 compared to mitosis might be due to the different processivities by which cohesin and condensin extrude DNA loops. Loop-forming cohesin is released from interphase chromatin after about 10-20 min by the protein WAPL (Gerlich *et al*, 2006b; Tedeschi *et al*, 2013), limiting DNA loop size (Haarhuis *et al*, 2017; Gassler *et al*, 2017; Wutz *et al*, 2017). In contrast, condensin II associates with mitotic chromatin more stably (Gerlich *et al*, 2006a; Walther *et al*, 2018) to form DNA loops that are substantially larger than those of interphase cells (Naumova *et al*, 2013; Gibcus *et al*, 2018). To test if cohesin’s capacity to resolve sister chromatids in G2 can be further tuned to promote more extensive sister chromatid resolution, we therefore searched for ways to specifically increase cohesin-mediated DNA looping.

The size of DNA loops formed by cohesin during interphase can be increased by depleting WAPL (Haarhuis *et al*, 2017; Gassler *et al*, 2017; Wutz *et al*, 2017), which also relocalises cohesin to axial structures on chromatin (Tedeschi *et al*, 2013). By inhibiting cohesin release, WAPL depletion leads to an increase in the amount of chromatin-associated cohesin, which could also potentially increase the amount of cohesive cohesin if the depletion is induced before cells complete DNA replication. Prior analyses of cells depleted of WAPL by RNAi reported a lower degree of sister locus separation in interphase (Eykelenboom *et al*, 2019) and less separation of whole sister chromatids in mitosis (Kueng *et al*, 2006; Gandhi *et al*, 2006; Nagasaka *et al*, 2016; Eykelenboom *et al*, 2019), but the extent to which these phenotypes are caused by effects on cohesion or loop extrusion has remained unclear.

To investigate more specifically how increasing cohesin-mediated DNA looping affects the global resolution of sister chromatids in G2 cells, we adapted our synchronisation regime such that we could selectively deplete WAPL in G2 (Fig. EV. 4a). Using this synchronisation procedure, we degraded WAPL homozygously tagged with FKBP12^F36V^ (Nabet *et al*, 2018) (here on referred to as dTAG) after cells had completed DNA replication (Fig. EV. 4b-d), thereby excluding potential effects of WAPL depletion on cohesion establishment, which can only occur during DNA replication (Uhlmann & Nasmyth, 1998). WAPL depletion in G2 cells increased the resolution of sister chromatids compared to control cells, while it also reorganised sister chromatids into parallel threads (Fig. 2a, b, d). Co-depletion of AID-tagged SMC4 together with dTAG-WAPL did not affect this organisation (Fig. EV. 4e-i, 5a-c), demonstrating that these structures form due to the activity of cohesin-mediated looping, independently of condensin. Furthermore, immunofluorescence-based detection of H3-S10 phosphorylation confirmed the G2 state of WAPL depleted cells with highly resolved sister chromatids (Fig. EV. 5d, e). Thus, when its loop-extruding processivity is increased, cohesin has the remarkable ability to resolve interphase sister chromatids to an extent that normally only occurs during mitosis – in the absence of condensins and other mitotic activities.

**Figure 2.**
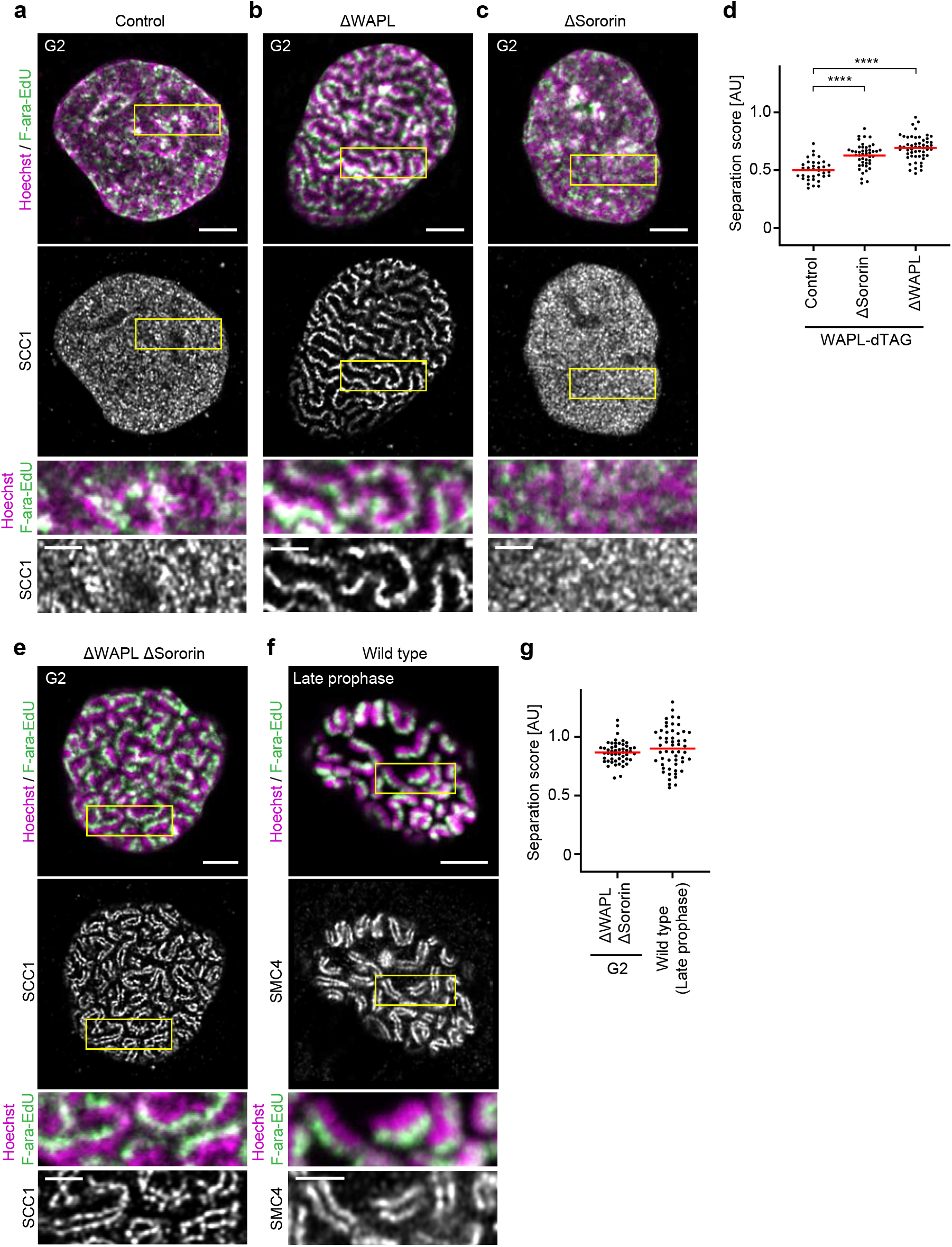
Hyperactive cohesin forms mitosis-like sister chromatids in G2. **a-c, e, f**, One sister chromatid was labelled per chromosome as in Fig. 1 and SMC4 (late prophase cells) or SCC1 (WAPL-dTAG cells) was visualised by immunofluorescence. **a**, Representative images of control WAPL-dTAG cells synchronised to G2 by RO-3306. **b**, Representative images of ΔWAPL cells synchronised to G2 by RO-3306. **c**, Representative images of ΔSororin cells synchronised to G2 by RO-3306. **d**, Quantification of sister chromatid separation as in Fig. 1d, e. Control WAPL-dTAG G2 (n = 36 cells from 6 experimental replicates), ΔSororin G2 (n = 45 cells from 6 experimental replicates) and ΔWAPL G2 (n = 53 cells from 6 experimental replicates) cells were analysed. Dots represent individual cells; red bars indicate the mean. Significance was tested using a two-tailed Mann Whitney U test; P = 5.90 × 10^−7^ (ΔSororin G2); P = 3.89 × 10^−11^ (ΔWAPL G2). **e**, Representative images of ΔWAPL ΔSororin cells synchronised to G2 by RO-3306. **f**, Representative images of wild type cells synchronised to late prophase by release from a RO-3306-mediated G2 arrest. **g**, Quantification of sister chromatid separation as in Fig. 1d, e. ΔWAPL ΔSororin G2 cells (n = 51 cells from 6 experimental replicates) and wild type late prophase (n = 55 cells from 7 experimental replicates) cells were analysed. Dots represent individual cells; red bars indicate the mean. All images show single Z-slices from 3D-stacks. Yellow boxes indicate inset regions. To aid visualisation, the SCC1 channel is not contrast matched. Scale bars large panels: 5 µm, insets: 2 µm.

Polymer simulations suggest that SMC-mediated loop extrusion organises DNA into a bottlebrush-like structure around a central axis of SMC protein complexes. Repulsion between the two bottlebrushes of sister chromatids then promotes their resolution (Goloborodko *et al*, 2016a). Removal of cohesive linkages between sister chromatid arms is essential for efficient resolution to occur in such simulations (Goloborodko *et al*, 2016a). We therefore wondered how cohesin can resolve sister chromatids so extensively in WAPL depleted G2 cells, which are expected to contain abundant arm cohesion. By visualising cohesin’s core subunit SCC1 using immunofluorescence, we observed a diffuse localisation of cohesin in control G2 cells, whereas after depletion of WAPL, cohesin localised to a single axis (Fig. 2a, b). The F-ara-EdU labelled chromatids consistently formed extended threads of DNA that localised adjacent to this cohesin axis, without intermixing across the axis with the sister chromatid not labelled by F-ara-EdU. These threads were often multiple microns in length, suggesting that adjacent DNA loops formed by cohesin consistently orient into the same direction over large genomic distances. Thus, in WAPL depleted cells, cohesin extensively resolves sister DNAs from a central axial location rather than forming two separate bottlebrush-like structures.

Reduced cohesion between sister chromatids might itself also function as a mechanism that promotes sister chromatid separation. To test how loss of cohesion affects the resolution of sister chromatids, we depleted Sororin, a protein required for the establishment and maintenance of cohesion (Rankin *et al*, 2005; Schmitz *et al*, 2007; Ladurner *et al*, 2016) but not for DNA looping (Davidson *et al*, 2019; Kim *et al*, 2019; Mitter *et al*, 2020) using RNAi (Fig. EV. 5f). In the absence of Sororin, sister chromatid resolution was increased compared to control cells, but not to the extent seen in WAPL depleted cells, and sister chromatids did not form extended threads (Fig. 2c, d). Thus, our data suggest that increased DNA looping processivity is a more powerful mechanism to resolve sister chromatids than removal of cohesion.

Cohesin’s localisation at the interface between sister chromatids in WAPL depleted G2 cells might result from steric constraints imposed by the cohesive pool of cohesin. To assess how cohesion affects the organisation of loop-forming cohesin, we let Sororin depleted cells replicate in the presence of WAPL such that cohesion between sister DNAs was lost, before synchronising cells to G2 and degrading WAPL. This perturbation indeed reorganised cohesin into pairs of separate axes (Fig. 2e), analogous to the organisation of condensins on late prophase chromosomes from wild type cells (Fig. 2f). Moreover, it resulted in a further increase in resolution compared to WAPL depletion alone, reaching levels of separation observed in late prophase (Fig. 2e-g). Thus, although chromosome arm cohesion prevents the separation of cohesin axes, it imposes only relatively small constraints on the resolution of sister chromatid DNA.

To investigate sister chromatid organisation in more detail relative to SMC protein axes, we analysed the distribution of DNA and SMC protein complexes along line profiles oriented perpendicular to the long axis of chromosomes. Measuring the distance between the peaks of F-ara-EdU-labelled and unlabelled DNA showed that the sister chromatids of WAPL depleted G2 cells were resolved almost as far apart as in unperturbed prophase or WAPL/Sororin depleted G2 cells (Fig. 3a-d), consistent with our separation score analysis. To assess the distribution of DNA relative to the SMC axes, we measured the distance between F-ara-EdU peaks and the closest SMC axis. This showed an outward-facing displacement of sister DNA in WAPL depleted G2 cells, in contrast to a symmetrical organisation in unperturbed prophase cells or WAPL/Sororin depleted G2 cells (Fig. 3e). Thus, in the presence of arm cohesion, DNA loop extrusion promotes a highly asymmetric displacement of sister chromatid DNA away from a central cohesin axis.

**Figure 3.**
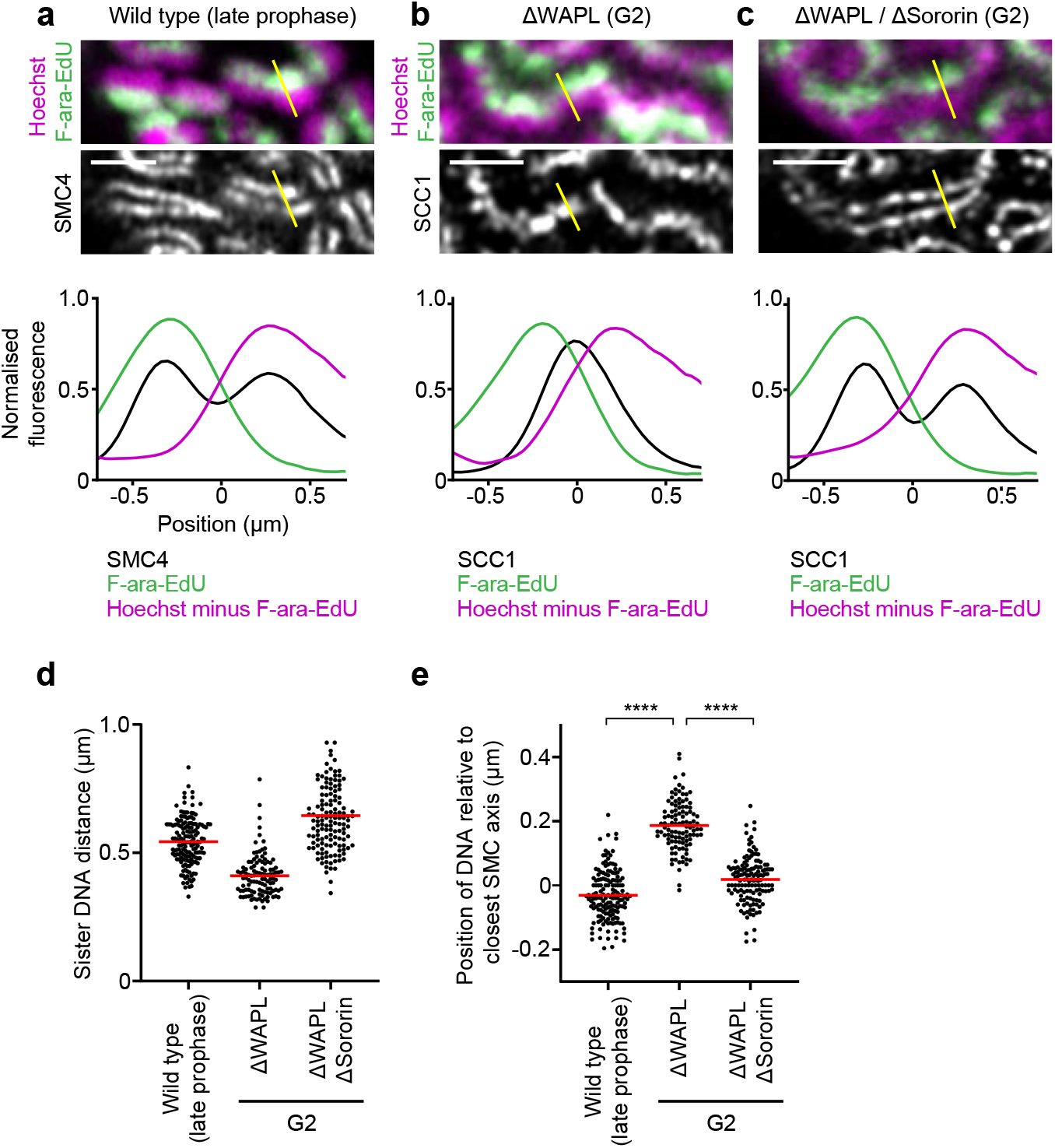
Organisation of sister chromatids around SMC protein axes. **a-c**, Analysis of sister DNA and SMC protein complex distribution in cross sections perpendicular to the long chromosome axis, as shown in Fig. 2b, e, f. Density of Hoechst minus F-ara-EdU was calculated by subtracting F-ara-EdU fluorescence from Hoechst fluorescence. Line profiles indicate the mean curves of individual line profile measurements aligned to the midpoint between two SMC4 peaks (wild type late prophase), the SCC1 peak (ΔWAPL G2 cells), or the midpoint between two SCC1 peaks (ΔWAPL ΔSororin G2 cells). **a**, Lines from wild type late prophase (n = 157 lines from 31 cells from 7 replicates). **b**, ΔWAPL G2 cells (n = 110 lines from 18 cells from 6 replicates). **c**, ΔWAPL ΔSororin G2 cells (n = 138 lines from 18 cells from 5 replicates) were analysed. **d**, Measurement of the sister DNA peak-to-peak distance for line profiles as in **a-c**. Dots represent individual distance measurements between chromatids; red bars indicate the mean. **e**, Measurement of the position of the F-ara-EdU labelled sister chromatid relative to the closest SMC peak for line profiles as in **a-c**. A positive value indicates an outward displacement relative to the axis, a negative value an inward displacement. Dots represent single line profile measurements; red bars indicate the mean. Significance was tested using a two-tailed Mann Whitney U test; P = 5.64 × 10^−40^ (late prophase); P = 3.52 × 10^−34^ (ΔWAPL ΔSororin G2). Yellow boxes indicate inset regions. Scale bars large panels: 5 µm, insets: 2 µm.

### Genomic range of sister chromatid resolution

The segregation of entire sister chromatids during mitosis requires DNA resolution over very large genomic distances. Imaging-based approaches based on sister-chromatid-specific fluorescence labelling as described above do not inform on the genomic ranges of sister chromatid resolution, and previous visualisation of pairs of neighboring genomic loci (Stanyte *et al*, 2018; Eykelenboom *et al*, 2019) also probe sister chromatid resolution only at a single local regime. To directly measure the genomic distance over which cohesin and condensin resolve sister chromatids, we therefore developed a new quantitative assay based on sister-chromatid-sensitive Hi-C (scsHi-C), a chromosome conformation technique that detects both intra- and inter-sister chromatid contacts based on labelling with the nucleotide analogue 4-thio-thymidine (Mitter *et al*, 2020, 2022).

In a closely juxtaposed and intertwined sister chromatid arrangement, the probability of a genomic locus contacting another genomic locus on the same DNA (cis sister contact) is expected to be as likely as contacting the sister DNA (trans sister contact), even over short genomic distances. With increasing degrees of sister chromatid resolution, cis sister contacts are expected to dominate trans sister contacts over increasing genomic intervals. On this basis, we calculated average cis/trans sister contact ratios over variable genomic distances and determined the interval at which cis sister contacts became as likely as trans sister contacts (considering a threshold slightly above noise), to derive a genomic resolution score (Fig. 4a).

**Figure 4.**
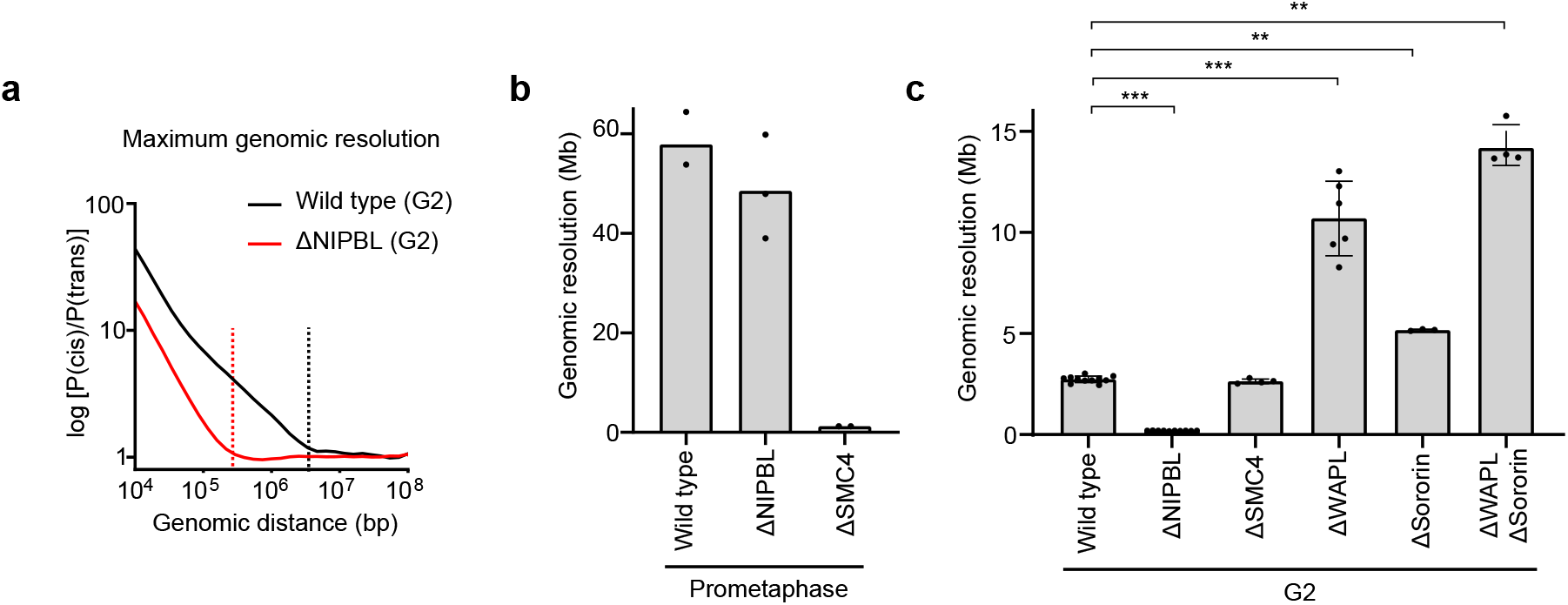
Genomic range of sister chromatid resolution. **a**, Analysis of sister chromatid resolution by sister-chromatid-sensitive Hi-C. Average contact probability curves over a range of genomic distances were calculated separately for cis and trans sister contacts (Extended Data Fig. 6) to derive cis/trans sister contact ratio curves for wild type (black) and ΔNIPBL (red) cells synchronised to G2. The genomic resolution intervals (dashed lines) were calculated by determining the genomic distance at which cis and trans sister contacts were equally abundant (at a threshold slightly above noise). **b**, Genomic resolution analysis for prometaphase cells. Wild type (n = 2 biologically independent experiments), ΔNIPBL (n = 3 biologically independent experiments) and ΔSMC4 (n = 2 biologically independent experiments) cells were analysed. Points indicate the values calculated for each replicate; bars indicate the mean. **c**, Genomic resolution analysis for G2 cells. Wild type (n = 11 biologically independent experiments), ΔNIPBL (n = 10 biologically independent experiments), ΔSMC4 (n = 4 biologically independent experiments), ΔWAPL (n = 6 biologically independent experiments), ΔSororin (n = 3 biologically independent experiments) and ΔWAPL ΔSororin (n = 4 biologically independent experiments) cells were analysed. The wild type and ΔSororin datasets were previously published in ^37^. Points indicate the values calculated for each replicate, bars indicate the mean and error bars indicate the standard deviation. Significance was tested using a two-tailed Mann Whitney U test; P = 1.24 × 10^−4^ (ΔNIPBL G2); P = 1.62 × 10^−4^ (ΔWAPL G2); P = 5.49 × 10^−3^ (ΔSororin G2), P = 1.47 × 10^−3^ (ΔWAPL ΔSororin G2).

With this genomic resolution assay, we analysed how cohesin and condensin contribute to sister chromatid resolution in interphase and mitosis (Fig. EV. 6, 7). In wild type prometaphase cells sister chromatids were resolved on average to 56.9 Mb (Fig. 4b; Fig. EV. 6a, d; Fig. EV. 7a). Mitotic sister chromatid resolution was almost completely suppressed by SMC4 depletion, whereas NIPBL depletion had little effect (Fig. 4b; Fig. EV. 6b-d; 7b-d), corroborating condensin’s key function in promoting long-range sister chromatid resolution in mitosis.

To study genomic resolution of sister chromatids in G2, we first analysed published scsHi-C data from wild type cells (Mitter *et al*, 2020), finding that their sister chromatids were resolved over 2.7±0.2 Mb (Fig. 4c; Fig. EV. 6e, h). We then performed scsHi-C in G2 synchronised cells depleted of NIPBL, detecting a more than 10-fold reduction in genomic resolution, whereas SMC4 depletion had no detectable effect (Fig. 4c; Fig. EV. 6f-h; 7e-h). These data confirm that the local resolution of sister chromatids in G2 depends on cohesin’s loop-extruding activity but not that of condensin.

To investigate how hyperactivating cohesin’s DNA loop-extrusion processivity affects sister chromatid resolution, we performed scsHi-C in G2 synchronised cells depleted of WAPL (Extended Fig. 6i, l; 7i-o). WAPL depletion increased the genomic resolution almost 4-fold over wild type cells, to 10.7±1.8 Mb, and resolution was further increased by co-depleting Sororin (Fig. 4c). However, depletion of Sororin alone had a smaller effect than WAPL depletion in the presence of Sororin, with an approximate 2-fold increase in maximum genomic resolution in Sororin depleted cells compared to wild type (Fig. 4c; Fig. EV. 6j-l). Together, our results show that removing cohesive linkages alone promotes the local separation of sister chromatids but is not sufficient to resolve sister chromatids over large genomic distances. In contrast, increasing the DNA loop-extruding processivity of cohesin is sufficient to resolve sister chromatids over large genomic distances, even in the absence of mitotic activities, supporting that chromatin loop extrusion is a fundamental mechanism by which genomes are segregated.

## Discussion

In our study, we have implemented novel assays for the detection of sister chromatid resolution based on microscopy and chromosome conformation capture. Our improved structural preservation and increased imaging resolution compared to previous work (Nagasaka *et al*, 2016) allowed us to study the fine structure of sister chromatid resolution already in interphase cells, whereas our scsHi-C assays enable direct measurement of the genomic distance over which sister chromatids resolve. In combination, these assays provide evidence for a previously underappreciated role of cohesin in sister chromatid organisation: cohesin not only holds sister chromatids together as previously known (Guacci *et al*, 1997; Michaelis *et al*, 1997; Losada *et al*, 1998; Gruber *et al*, 2003; Ivanov & Nasmyth, 2005; Watrin *et al*, 2006; Haering *et al*, 2008), but also moves them apart, most likely via its DNA loop-extruding activity, resulting in a balance between opposing forces (Fig. 5a).

**Figure 5.**
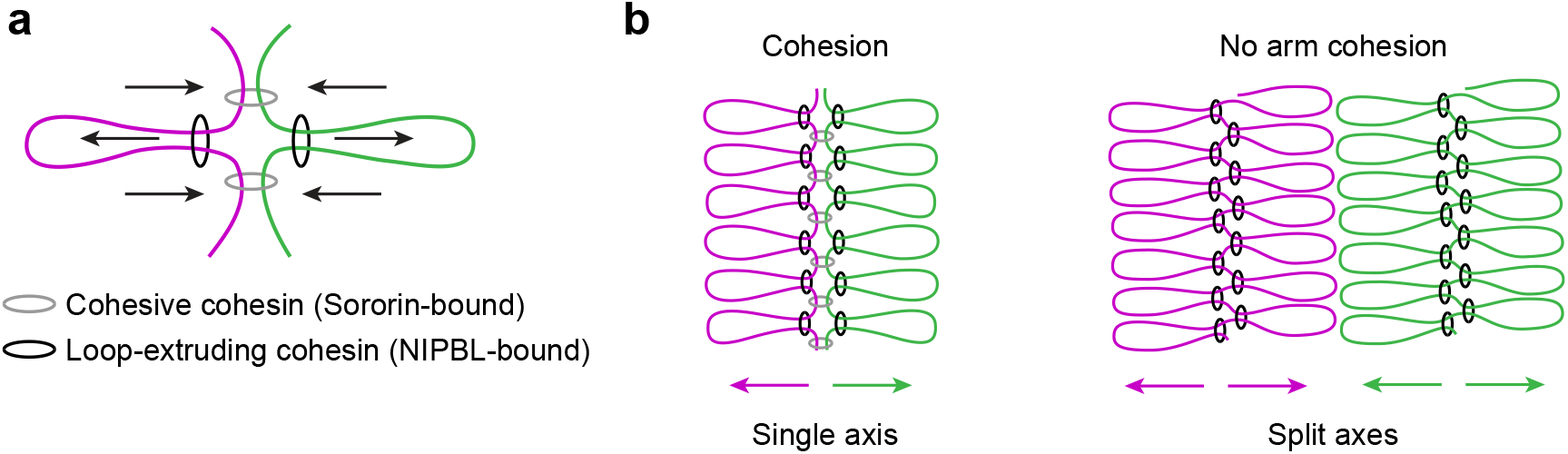
Models for sister chromatid organisation by cohesin. **a**, Cohesin associated with NIPBL extrudes DNA loops to move sister chromatids apart, whereas cohesin associated with Sororin maintains linkages between sister chromatids. **b**, Hyperactivating cohesin’s loop extrusion processivity resolves sister chromatids in G2 despite the presence of arm cohesion, resulting in an asymmetrical distribution of adjacent DNA loops relative to a central cohesin axis. Additional removal of cohesion along chromosome arms promotes cohesin axis separation, resulting in a symmetrical organisation of DNA loops around two separate cohesin axes. Magenta and green indicate sister DNAs, grey and black rings indicate cohesive and loop-extruding cohesin, respectively.

Our finding that increasing cohesin’s looping capability by depleting WAPL induces prophase-like sister chromatid resolution is consistent with data from meiotic chromosomes where a meiosis specific variant of cohesin organises chromatids into threads (van Heemst *et al*, 1999; Klein *et al*, 1999; Zwettler *et al*, 2020; Ur & Corbett, 2021) and also with previous polymer simulations, which showed that a transition from a scarce regime with gaps between loops to a dense regime without gaps promotes efficient resolution of sister DNAs (Goloborodko *et al*, 2016a, 2016b). Counter to model predictions (Goloborodko *et al*, 2016a), however, our data show that sister chromatids can resolve their DNA over large genomic intervals even in the presence of abundant arm cohesion. In WAPL depleted cells, sister DNAs resolve around a single cohesin axis that presumably contains both loop-forming and cohesive cohesin. Such resolution in the presence of cohesion might be driven by steric repulsion between DNA loops (Polovnikov *et al*, 2022) when the directionality of loop extrusion is coordinated between neighboring loops (Fig. 5b).

Prior work suggested that sister chromatid resolution is impaired in WAPL-depleted cells (Kueng *et al*, 2006; Gandhi *et al*, 2006; Nagasaka *et al*, 2016; Eykelenboom *et al*, 2019). Our improved assays and time-controlled depletion, however, clarify that when WAPL is depleted after completion of DNA replication, it strongly promotes sister DNA resolution rather than impairing it. As the sister DNAs organise around a single axis of cohesin, however, their resolution cannot be detected by conventional bulk DNA staining as in prior work (Kueng *et al*, 2006; Gandhi *et al*, 2006). Moreover, the phenotype analysis of RNAi-mediated WAPL depletion (Kueng *et al*, 2006; Gandhi *et al*, 2006; Nagasaka *et al*, 2016; Eykelenboom *et al*, 2019) is inherently limited by potential effects on both cohesin-mediated DNA looping and cohesin establishment, and therefore difficult to interpret.

We provide evidence that the resolution of sister chromatids into individual bodies does not require mitosis-specific activities, apart from increased DNA loop-extrusion processivity, while reduced cohesion between sister chromatids facilitates the additional separation of SMC-protein axes. Thus, while changes in histone posttranslational modifications (Cimini *et al*, 2003; Wilkins *et al*, 2014; Zhiteneva *et al*, 2017; Schneider *et al*, 2022) and the coating of chromosomes with a repulsive surface layer (Cuylen *et al*, 2016) are essential for proper interactions with spindle microtubules (Schneider *et al*, 2022) and nuclear assembly (Samwer *et al*, 2017; Cuylen-Haering *et al*, 2020), the crucial initial step of pre-segregating sister chromatids can largely be mediated by increasing DNA loop extrusion processivity alone.

Although vertebrate cells strictly rely on condensin to resolve and segregate their sister chromatids in mitosis, budding yeast can to some extent resolve and segregate many chromosomal regions even in the absence of condensin, except for repetitive rDNA loci (Freeman *et al*, 2000). This might be explained by the functional equivalence of cohesin- and condensin-mediated sister chromatid resolution by DNA loop extrusion, as demonstrated in our study, given that in budding yeast cohesin is not removed from chromosome arms during mitotic entry. In contrast to nucleosomes, SMC protein complexes contributing to chromosome segregation are also present in archaea and bacteria (Wang *et al*, 2015; Palecek & Gruber, 2015; Wang *et al*, 2017). Thus, DNA loop extrusion might be the most ancient and universal genome-segregating activity shared across all kingdoms of life.

## Materials and Methods

### Cell culture

Each cell line used in this study has regularly tested negative for mycoplasm contamination. All cell lines in this study are derived from a parental HeLa cell line (Kyoto strain), which was validated using a Multiplex Human Cell line Authentication test (MCA) or from a HeLa Kyoto cell line previously described in (Schmitz *et al*, 2010). Cells were grown in Dulbecco’s modified eagle medium (DMEM, IMP/IMBA/GMI Molecular Biology Service/Media Kitchen), supplemented with 10% (v/v) fetal bovine serum (FBS): (Gibco, 10270-106, 2078432), 1% (v/v) penicillin-streptomycin (Pen/Strep): (Sigma Aldrich, P0781), GlutaMAX: (Invitrogen, 35050038), (here on referred to as wild type medium), in combination with antibiotics, depending on the resistance of the respective cell line. To culture HeLa SMC4-AID, HeLa NIPBL-AID and WAPL-dTAG cell lines, cells were supplemented with blasticidin S (6 µg mL-1, Thermo Fisher Scientific, A1113903). The WAPL-AID cell line was supplemented with puromycin (0.5 µg mL-1, Calbiochem, 540411). HeLa SMC4-AID_WAPL-dTAG cells were supplemented with blasticidin S (6 µg mL-1, Thermo Fisher Scientific, A1113903) and hygromycin B (0.3 mg mL-1, Roche, 10843555001). All cell lines were cultured in a humidified incubator at 37°C, with 5% CO_2_.

### Genome editing using CRISPR/Cas9

The SMC4-AID cell line used in this study was generated in (Schneider *et al*, 2022). The NIPBL-AID cell line was generated using a chimeric Cas9-human geminin fusion (Cas9-hGem), originally published in (Gutschner *et al*, 2016). A single guide RNA (sgRNA) (sequence: caccgtgtccccattactactcttg) was cloned into the Cas9-hGem expressing plasmid. To endogenously tag NIPBL at its N-terminus with a mini AID degron (mAID) (Kubota *et al*, 2013), a repair template was designed with homology arms of 682 bp and 685 bp flanking the insertion site 5’ and 3’ respectively. The repair template contained the following elements: 5’ homology arm, mEGFP tag for visualisation, 2 x Gly-Gly-Ser linker, mAID, Mattaj-Tandem linker, 3’ homology arm. The protospacer adjacent motif (PAM) was also mutated to prevent re-editing of successfully edited clones. The repair template was produced as a gBlock® (IDT) and cloned into a donor vector (pCR2.1) for amplification. The sgRNA/Cas9 and repair template containing plasmids were co-transfected into cells using the Neon™transfection system, using the standard manufacturer protocol for HeLa cells, as described in (Schneider *et al*, 2022). 9 days after transfection, mEGFP positive clones were sorted into 96-well plates. Homozygously edited clones were identified by genotyping as described in (Samwer *et al*, 2017) using the following primers: gcctagcagttaagaaacaaact (forward), gctccaacagattactgaacatga (reverse). The OsTIR1^F74G^ ligase (Yesbolatova *et al*, 2020) was stably integrated into a homozygously edited clone at the ‘safe harbour’ adeno-associated virus integration (AAVS1) site as described previously (Li *et al*, 2019; Schneider *et al*, 2022). Clones which efficiently depleted NIPBL after integration of the ligase were identified by treating clones with 1 µM 5-Ph-IAA (BioAcademia, 30-003) for 90 min, before analysis by flow cytometry using an iQue Screener Plus instrument. The WAPL-dTAG cell line was generated using a nickase Cas9 strategy originally described in (Ran *et al*, 2013). The guide RNA and repair template plasmids were as described in (Nagasaka *et al*, 2022). In brief, two sgRNAs were cloned into separate nickase Cas9 expressing plasmids (guide1 sequence : caccgctaagggtagtccgtttgt, guide2 sequence : caccgtggggagagaccacattta). To endogenously tag WAPL at its N-terminus, a repair template was designed with homology arms of 1108 and 719 bp flanking the insertion site 5’ and 3’ respectively. The repair template contained the following elements: 5’ homology arm, blasticidin resistance cassette, Gly-Ser-Gly linker, P2A sequence, 2 x HA tag, Gly-Ser-Gly linker, FKBP12^F36V^ (dTAG) sequence, 15 x Gly linker, 3’ homology arm. The repair template was cloned into a donor plasmid (pBSKII) for amplification. The sgRNA/Cas9 and repair template containing plasmids were also co-transfected into cells using the Neon™transfection system. 8 days after transfection antibiotic selection was performed using 6 µg mL-1 blasticidin S (Thermo Fisher Scientific, A1113903) to identify cells which had undergone productive editing. Homozygously edited clones were identified by genotyping using the following primers: gcacaaagctctcttggcggag (forward), gtcacagcgcaaattacattacaccaag (reverse). Cells which efficiently depleted WAPL were identified using confocal microscopy and Western blotting after treatment of the cells with 1 µM dTAG-7 (Tocris, 6912) for 3 h. To generate the SMC4-AID_WAPL-dTAG cell line, WAPL was endogenously tagged at its N-terminus with dTAG in the SMC4-AID cell line referenced above and published in (Schneider *et al*, 2022). The same sgRNAs were used as described for the WAPL-dTAG cell line. A different repair template was used, with homology arms of 700 bp either side of the site of insertion. The repair template contained the following elements: 5’ homology arm, hygromycin resistance cassette, Gly-Ser-Gly linker, P2A sequence, 2 x HA tag, 5 x Gly linker, FKBP12^F36V^(dTAG), Mattaj Tandem linker, 3’ homology arm. The plasmids were co-transfected into the cells using electroporation as described above. 8 days after transfection, productively edited cells were selected with 0.3 mg mL-1 hygromycin B (Roche, 10843555001). Homozygously edited clones were identified using the same genotyping strategy as described above and cells which efficiently depleted WAPL were identified using confocal microscopy and Western blotting. The Halo-AID-WAPL_SCC1-mEGFP cell line was generated by first endogenously tagging SCC1 at its C-terminus with monomeric EGFP (L221K) using the same guides and genotyping primers described in (Wutz *et al*, 2017). WAPL was then N-terminally tagged with a mini-AID (amino acids (aa) 71-114) degron (Morawska & Ulrich, 2013), using the same gRNAs described above. The repair template contained homology arms of 1108 and 719 bp 5’ and 3’ to the insertion site respectively and contained the following elements: 5’ homology arm, Halo tag, Ala-Ser-Gly-Leu-Arg-Ser-Arg-Gly linker, mAID, 5 x Gly linker, 3’ homology arm. Homozygously edited clones were identified using the following genotyping primers : tgatttttcattccttaggcccttg (forward), tacaagttgatactggccccaa (reverse). OsTIR1 was introduced into a homozygously edited clone using lentivirus as described in (Wutz *et al*, 2017) and cells were characterised as described above.

### Sister chromatid labelling cell culture protocol

All cell lines were synchronised using a four block and release protocol, apart from NIPBL-AID cells, where a three block and release protocol was used instead. For a four block and release protocol, cells were first pre-synchronised to the G1/S boundary by incubation with 2 mM thymidine (Sigma Aldrich, T1895) in wild type medium for 16-18 h. Cells were then washed twice with pre-warmed PBS and released into fresh medium. 10-12 h after release, cells were blocked again using 3 µg mL-1 aphidicolin (Sigma Aldrich, A0781). To generate one-sister chromatid labelled cells, 10 µM F-ara-EdU (Sigma Aldrich, T511293) was added to cells 15 h after the second release into S phase (after the first release for the NIPBL-AID cell line) to generate a pool of the compound within the cell, while the cells were in aphidicolin. The following day, cells were washed twice with pre-warmed PBS and released into fresh medium containing 10 µM F-ara-EdU. A final block with aphidicolin was then performed as described above. The following day, cells were washed twice with pre-warmed PBS and released into fresh nucleotide-free medium, such that the cells became labelled on only one sister chromatid. To generate sister chromatids labelled on two chromatids, 10 µM F-ara-EdU was added at the time of the final aphidicolin block. For the final S phase release, cells were then released into medium containing 10 µM F-ara-EdU.

### Sister chromatid labelling – G2 and mitotic arrests, and protein depletions

After labelling of one or two sister chromatids, cells were synchronised to G2 using the CDK1 inhibitor RO-3306 (Sigma Aldrich, SML0569). RO-3306 was added 5-6 h after the release from the final aphidicolin block. Protein depletions, siRNA transfections and the timing of fixation were performed as follows: Wild type G2 samples were fixed 15 h after the release from the final aphidicolin block. To deplete SMC4 in the SMC4-AID cell line, 1 µM 5-Ph-IAA (BioAcademia, 30-003) was added 1 h before the release from the final aphidicolin block, and samples were fixed 15 h after the release. For NIPBL-AID experiments, 1µM 5-Ph-IAA was added 10 h after the release from the final (third) aphidicolin block, and cells were fixed 15 h after release. To deplete WAPL in WAPL-dTAG or SMC4-AID_WAPL-dTAG cell lines, 1 µM dTAG-7 (Tocris, 6912) was added 14 h after the release from the final aphidicolin block. To deplete SMC4 in the SMC4-AID_WAPL-dTAG cell line, 1 µM 5-Ph-IAA (BioAcademia, 30-003) was added 1 h before the release from the final aphidicolin block. For experiments using the WAPL-dTAG or SMC4-AID_WAPL-dTAG cell lines, samples were fixed 24 h after the release from the final aphidicolin block, and siRNA transfections using XWNeg or Sororin siRNAs was performed as described below (section, siRNA transfection). G2 samples fixed after 15 h were treated with 8 µM RO-3306, while G2 cells fixed after 24 h were treated with 10 µM RO-3306. To generate prometaphase samples, 7 µM RO-3306 was added to cells as described above. 15 h after release from the final aphidicolin block, cells were washed three times with pre-warmed wild type medium. After the final wash, cells were released into wild type medium containing 5 µM STLC (Enzo Life Sciences, ALX-105-011-M500), and fixed 30 - 60 minutes later. For all samples where proteins were depleted, 5-Ph-IAA or dTAG-7 were added to all wash buffers and medium after they had been added to the samples.

### Immunofluorescence

Immunofluorescence was performed in parallel in all sister chromatid labelling experiments, before click chemistry was performed. In all experiments, cells were grown on Nunc™LabTek™8-well chambered cover glass (8-well Labteks). At the time of fixation, cells were washed twice with PBS before fixation with PBS containing 4% methanol-free formaldehyde (Thermo Fisher Scientific, 28906) for 5 min. The fixative was quenched using 10 mM TRIS-HCl (Sigma Aldrich) pH 7.5 in PBS for 3 min. Cells were permeabilised by incubation with PBS containing 0.2% Triton-X-100 (Sigma Aldrich, X100-100 mL) for 5 min, washed once with PBS to remove residual detergent, and then blocked for 30 min with 0.45 µm filtered 2% BSA (Sigma-Aldrich, A7030) in PBS (blocking buffer). Primary and secondary antibodies were diluted with blocking buffer. All primary antibody incubations were performed for > 16 h at 4 °C with gentle shaking. Incubations with the anti-GFP nanobody were performed at room temperature for 2 h. After primary incubations, samples were washed 3 × 10 min with PBS, before incubation with secondary antibodies for 1 h at room temperature with gentle shaking. Samples were then washed 3 × 10 min with PBS. For samples labelled with F-ara-EdU, click chemistry staining was performed as described below (section: click chemistry). Otherwise, Hoechst 33342 was added to a final concentration of 1.62 µM in the wash buffer after secondary antibody incubation. Proteins were visualised using the following primary and secondary antibody combinations: Cyclin B1 was detected with a monoclonal rabbit antibody (Cell Signaling Technology, 12231S, 1:800) and visualised using a goat anti-rabbit Alexa Fluor 647 secondary antibody (Molecular Probes, A21244, 1:1000) or detected with a monoclonal mouse antibody (SantaCruz, sc-245, 1:250) and visualised with a donkey anti-mouse Alexa Fluor 647 secondary antibody (Molecular Probes, A31571, 1:1000). SCC1/ RAD21 was detected with a monoclonal mouse antibody (Millipore, 05-908, 1:500) and visualised with a goat anti-mouse Alexa Fluor 568 secondary antibody (Molecular Probes, A11004, 1:1000) or a goat anti-mouse Alexa Fluor 488 secondary antibody (Molecular Probes, A11001: 1:1000). SMC4 was detected using a monoclonal rabbit antibody (Abcam, ab229213, 1:250) and visualised with a goat anti-rabbit Alexa Fluor 568 secondary antibody (Molecular Probes, A11011, 1:1000). HA-WAPL was detected using a monoclonal rabbit antibody against the HA-tag (Cell Signalling Technology, 3724S, 1:1000) and visualised with a goat anti-rabbit Alexa Fluor 568 secondary antibody (Molecular Probes, A11011, 1:1000). Phosphorylated Histone H3-Ser10 was detected using a monoclonal mouse antibody (Millipore, 05-806, 1:5000) and visualised using a goat anti-mouse Alexa Fluor 568 secondary antibody (Molecular Probes, A11004, 1:1000). mEGFP-mAID-NIPBL and SCC1-mEGFP were detected and visualised with an anti-GFP nanobody booster (Chromotek, gba488-100, 1:250).

### Click chemistry

All immunofluorescence staining was performed before the click chemistry reaction. Visualisation of F-ara-EdU labelled sister chromatids was performed using Molecular Probes Click iT Cell Reaction Buffer Kit (ThermoFisher, C10269), in combination with AF-488-Picoyl-Azide (Jena Bioscience, CLK-1276-1). 10x Click iT reaction buffer was diluted 1 in 10 with monoQ H_2_O to make a 1 x solution freshly before use. A 1 x solution of Click iT buffer additive was prepared with the same diluent freshly before use. The click reaction cocktail was prepared by mixing the following components (in the same order as they are written here) according to manufacturer instructions: 1 x Click-iT reaction buffer, AF-488-Picolyl-Azide (5 µ M final concentration), copper sulphate (2 mM final), 1 x Click-iT buffer additive. The reaction cocktail was added to cells for 30 min with gentle rocking on a shaker, protected from light. The cocktail was removed, and the cells incubated for 5 min with 0.45 µm filtered 2% BSA in PBS. Cells were then washed twice with 1.62 µM Hoechst 33342 for 3 × 10 min to stain DNA, and then stored at 4 °C until imaging.

### siRNA transfection

siRNA transfections using Sororin and XWNeg siRNAs were performed in 8-well Labteks. siRNAs were transfected using lipofectamine RNAiMax (Life Technologies, 13778150) according to the manufacturer instructions. Sororin was targeting using a 16 nM custom silencer select siRNA originally published i n (Schmitz *et al*, 2007), (sensestr and, GCCUAGGUGUCCUUGAGCUtt, Ambion, including a 3’ tt overhang for increased efficiency). XWNeg was used as a custom silence select siRNA at 16nM (sense strand, UACGACCGGUCUAUCGUAGtt, Ambion, including a 3’ tt overhang for increased efficiency). For microscopy experiments, the transfection mix was added to cells 6 h after the third release (see above, sister chromatid labelling for details). The transfection mix was left on the cells for 16-18 h, before being washed out and replaced with wild type medium. For scsHi-C experiments, siRNA transfections using the same siRNAs were performed as described, but in 25 cm^2^ flasks, and using 32 nM siRNAs.

### Labelling cells with Halo-TMR

SMC4 was stained using HaloTag TMR Ligand (Promega, G8251) in control and ΔSMC4 cells for 30 min (1:1000 dilution in wild type medium). Cells were then washed three times with fresh wild type medium. The medium was exchanged once more, and cells left for 30 min in the absence of ligand before fixation as described above (section: Immunofluorescence).

### Microscopy and image processing

All sister chromatid labelling experiments were imaged on a custom Zeiss LSM 980 microscope fitted with an additional Airyscan2 detector, using a 63 x NA 1.4 oil DIC Plan-Aphrochromat (Zeiss) objective and ZEN 3.3 Blue 2020 software. Experiments were performed with optimal sectioning (150 nm between each Z-slice) using the AiryScan SR (Super Resolution) mode. 3D Airyscan processing was performed for each acquired image using default parameters. Airyscan processed images were then registered to correct for chromatic shift from the microscope using either TetraSpeck™ Fluorescent Microspheres Size Kit (mounted on slide) (Thermo Fisher Scientific, T14792, 0.2 µm bead size) or TetraSpeck™ Microspheres, 0.2 µm (Thermo Fisher Scientific, T7280, prepared in-house and spread onto glass slides) in combination with a custom Fiji(Schindelin *et al*, 2012) script. A four colour Z-stack image of the bead slide was acquired using the same Z-section interval and zoom as the images to be registered, before running the script to register the cells in X, Y and Z dimensions. As the AiryScan processing procedure added an additional 10000 grey pixel values to each processed image, 10000 pixel values were subtracted from each registered image in Fiji using the ‘Subtract’ module before quantification. Experiments to measure protein depletions were performed on a custom Zeiss LSM 780 microscope using a 63 x NA 1.4 oil DIC Plan-Aphrochromat (Zeiss) objective and ZEN Black 2011 software.

### Image analysis – separation score

The separation score for each condition was calculated using a custom pipeline written in Python. In brief, the pipeline performed the following steps. For each processed Z-stack image, the central slice was calculated by determining the centre of mass of the chromatin (Hoechst) channel, using the scipy ‘ndimage’ module (Virtanen *et al*, 2020). A subset of slices around the central slice (5 above, 5 below, 11 slices in total) was chosen for the F-ara-EdU and Hoechst channels for each image. For each chosen slice, the Hoechst channel was segmented using Otsu thresholding to generate a chromatin mask. By default, objects which touched the image border were excluded using the ‘clear_border’ functionality from the skimage ‘segmentation’ module (van der Walt *et al*, 2014). These cells were subsequently analysed separately with the ‘clear_border’ parameter switched off. Small objects and cellular debris were excluded from the mask with a size filter using the ‘remove_small_objects’ functionality from the skimage ‘morphology’ module (van der Walt *et al*, 2014). The chromatin mask was applied to the F-ara-EdU and Hoechst channels, and all pixel values within the mask were extracted. The Spearman correlation coefficient (SCC) between the two channels was then calculated to give a single SCC value per slice. This operation was performed for each chosen slice, before subsequently calculating the mean SCC per cell. The SCC value per cell for each condition was then normalised relative to the mean SCC values of wild type two (0 value) and one (1 value) sister labelled prometaphase cells, to generate the separation score metric.

### Image analysis – distribution of sister DNAs around SMC protein axes

Line profiles were drawn across single or split SMC protein axes in Fiji using the line profile tool with a line width of 5 pixels. All cells analysed were one-sister chromatid labelled. The line profiles were drawn across axes in a consistent way, moving always from the F-ara-EdU labelled chromatid to the unlabelled chromatid. The line profile coordinates and pixel intensities at each position were then extracted and saved as a csv file using a custom Fiji script. All line profile regions of interest (ROIs) from this analysis were also saved. The data was then further analysed using a custom Python script. The line profile distances and pixel intensities for the SCC1/SMC4, F-ara-EdU and Hoechst channels were extracted from the csv file. A min/max normalisation was performed on each channel to account for the differences in raw signal intensity. To overcome the limitation that only one sister chromatid was specifically labelled using F-ara-EdU in our protocol, the normalised F-ara-EdU signal was subtracted from the normalised Hoechst signal at positions where the normalised F-ara-EdU values were greater than the normalised Hoechst values. This subtraction operation generated a separate ‘Hoechst minus F-ara-EdU’ plot profile. The Hoechst minus F-ara-EdU profile (here on referred to simply as Hoechst) was then again min/max normalised to rescale the data. A polynomial fit was then performed on the normalised SCC1/ SMC4, F-ara-EdU and Hoechst profiles using the numpy ‘poly1d’ operation. Peaks were then identified for the three profiles using the scipy ‘signal’ module (Virtanen *et al*, 2020). For WAPL depleted cells, only profiles with a single peak for the SCC1, F-ara-EdU and Hoechst channels were considered for the downstream analysis. For all other conditions, where split axes are expected, only profiles with two peaks for the SCC1/SMC4 channel and a single peak for the F-ara-EdU and Hoechst channels were considered. The amount of sister chromatid resolution around single or split SMC protein axes was calculated by measuring the peak-to-peak distance of the normalised F-ara-EdU and Hoechst profiles. To calculate the radial displacement of sister chromatids relative to a single axis, the distance between the SCC1 peak and the F-ara-EdU peak was calculated. To calculate the radial displacement of sister chromatids relative to split SMC protein axes, the distance between the F-ara-EdU peak and the closest SCC1/SMC4 axis was calculated. To plot the data, normalised Hoechst and F-ara-EdU mean curves were aligned relative to single or split SMC protein axes. For resolved sister chromatids around a single cohesin axis, for each set of profiles the peak of the normalised SCC1 channel was identified (as described above) and the normalised F-ara-EdU and Hoechst channels aligned relative to this. For cells with split axes, the two peaks for the normalised SCC1 or SMC4 profile were identified and the midpoint between the two peaks calculated. The normalised Hoechst and F-ara-EdU profiles were then aligned relative to this midpoint.

### Image analysis – measurement of mean pixel intensities, single cells

To measure mean intensities of single cells, a custom Python script was used. For mean intensity measurements, the central slice of each Z-stack was first determined as described above, by identifying the centre of mass of the chromatin (Hoechst) channel using the scipy ‘ndimage’ module (Virtanen *et al*, 2020). Mean intensity measurements were then performed for each channel in this central slice. For all measurements, the Hoechst channel was segmented to create a mask. For mean intensity measurements, G2 and prophase cells were segmented using Li thresholding and prometaphase cells using Otsu thresholding. The mask was then applied to all channels of interest and mean fluorescence within the mask calculated. In Extended Data Fig. 2b, 5d, background fluorescence in the phospho-H3-Ser10 channel was calculated by measuring fluorescence in non-chromatin regions of five prometaphase cells and calculating the mean. The data were then normalised relative to the mean background fluorescence of the phospho-H3-Ser10 channel (0 value) and mean phospho-H3-Ser10 fluorescence of prometaphase (1 value) cells for the conditions of interest. In Extended Data Fig. 2d, the background fluorescence in the cyclin B1 channel was calculated by measuring fluorescence in non-cell regions of five G2 cells and calculating the mean. In Extended Data Fig. 2d, the data were normalised relative to the mean background fluorescence of the cyclin B1 channel (0 value) and the mean nuclear cyclin B1 fluorescence of prophase (1 value) cells.

### Image analysis – measurement of mean pixel intensities, fields of cells

Fields of cells were analysed using a custom Python script. A Gaussian blur was applied to the Hoechst channel (sigma = 1.0) using the skimage ‘filters’ module and the Hoechst channel was then segmented using Li thresholding. Cells which touched the border of the image were excluded using the ‘clear_border’ functionality from the skimage ‘segmentation’ module. Individual masks were labelled and applied to each cell in the field. Overlapping cells were distinguished using the ‘watershed’ functionality of the skimage ‘segmentation’ module. The area of the nuclear mask and mean fluorescence within the mask was then calculated. In Extended Data Fig. 3d, 4f, 7j, wild type cells were stained with Halo-TMR and the mean Halo-TMR fluorescence within the segmented nuclei then calculated. The data were then normalised relative to the mean nuclear Halo-TMR fluorescence of wild type cells (0 value) and control cells (1 value). In Extended Data Fig. 3h, 4d, 4g, background fluorescence in the channel of interest was calculated by measuring fluorescence in non-cell areas of three fields and calculating the mean. The data were then normalised relative to the mean background fluorescence of the channel of interest (0 value) and the mean nuclear fluorescence of the protein of interest in control cells (1 value).

### Image analysis – calculation of percentage labelled sister chromatids per cell in prometaphase cells

For wild type two and one-sister chromatid labelled prometaphase cells, sister chromatid pairs were identified using the Hoechst (to identify DNA) and SMC4 (to identify axes) channels. With only these two channels switched on, line profiles were drawn in Fiji (5-pixel line width) along the length of the chromatid pair. The length of each line for the Hoechst channel was then measured for each line in the cell, the ROIs saved, and the measurements saved as a csv file. For each measured line, the F-ara-EdU channel was then switched on, and the length of 0, 1 or 2 sister chromatid labelled fragments measured within the original measured line, and then saved as a csv file. A custom Python script was then used to sort and group the data such that the percentage of 0, 1 or 2-sister labelled fragments was calculated on a per cell basis.

### Image analysis – classification of late prophase cells

Cells were identified as late prophase through immunostaining to determine cyclin B1 and SMC4 mean intensity and localisation, in addition to staining with Hoechst 33342 to assess chromosome morphology. Mean nuclear cyclin B1 fluorescence was determined in a central Z-section as described above, and the cells classified into three classes (early, mid, late prophase) based on mean nuclear cyclin B1 intensity. After this stratification, cells were manually verified based on chromosome morphology, extent of SMC4 axis formation and localisation of cyclin B1.

### scsHi-C cell culture

Cell culture for G2 scsHi-C was performed as described in (Mitter *et al*, 2020, 2022). Wild type medium without selection antibiotics was used to synchronise the cells, and all chemicals used were diluted in this medium to the specified concentration. All centrifugation steps before formaldehyde fixation were performed at 1100 g, and all centrifugations after fixation at 2500 g. In brief, asynchronous cells were synchronised to the G1/S boundary with 2 mM thymidine (Sigma Aldrich, T1895) for 16-18 h. The following day, cells were washed twice with prewarmed PBS, before the addition of fresh wild type medium. 10-12 h post release, 3 µg mL-1 aphidicolin (Sigma Aldrich, A0781) was added to the cells to arrest them at the G1/S boundary. 2 mM 4-thiothymidine (4sT) (Carbosynth, NT06341) was also supplemented to the medium at this stage. 16-18 h hours later, cells were washed twice with prewarmed PBS before the addition of 2 mM 4sT. For all G2 samples shown in Fig. 4, 10 µM RO-3306 (Sigma Aldrich, SML0569) was added to the cells 5-6 h after the second release, and samples were harvested 24 h after the second release. Experiments with protein depletions were performed as follows: For SMC4-AID experiments, 5-Ph-IAA (BioAcademia, 30-003) was added 1 h before the second release to a final concentration of 1 µM. For NIPBL-AID experiments, 5-Ph-IAA was added 8 h after the second release to a final concentration of 1 µM. For WAPL-AID experiments, indole-3-acetic-acid (auxin) (Sigma Aldrich, I5148) was added to a final concentration of 500 µM 14 h after the second release. For all samples where proteins were depleted, auxin or 5-Ph-IAA were added to all wash buffers and medium after they had been added to the samples. For WAPL-AID experiments, siRNA transfections using XWNeg or Sororin siRNAs was performed as described above (section: siRNA transfection), 6-7 h after the release from the thymidine block. To harvest cells, cells were washed twice with PBS, with banging against a solid support each time to remove dead and mitotic cells. Cells were then trypsinised to detach them from the culture flask before quenching with wild type medium. Collected cells were spun down and the pellet washed once with PBS. Cells were spun down, PBS removed, and the cells were fixed with PBS containing 1% methanol-free formaldehyde (Thermo Fisher Scientific, 28906) for 4 min with gentle rotation. Cells were spun down, the formaldehyde removed and 20 mM TRIS-HCl pH 7.5 in PBS added to quench residual formaldehyde. Samples were spun down once more, the liquid removed, and samples stored at -20 °C until further processing. To generate prometaphase samples, cells were treated as described above until the addition of RO-3306. Cells were also cultured in 75 cm^2^ rather than 25 cm^2^ flasks to generate sufficient cell numbers for the experiment. RO-3306 was added to a final concentration of 7 µM to cells 7 h after the second release. Before adding RO-3306, cells were washed twice with PBS and banged against a solid support to remove dead and mitotic cells. 14 h after the second release, cells were washed three times with wild type medium. After the final wash, wild type medium containing 200 ng mL-1 nocodazole (Sigma Aldrich, M1404) was added to the cells. 60 minutes later, mitotic cells were detached from the culture flask by banging the flask against a solid support. The medium was then collected, cells spun down and washed once with PBS, before fixation with 1% formaldehyde as described above.

### Library prep and scsHi-C sample preparation

Hi-C sample preparation was performed as described in (Mitter *et al*, 2020, 2022). All centrifugations were performed at 2500 g, apart from those in the ethanol precipitation step, which were performed at 21000 g. Frozen cell pellets were thawed and then lysed using ice cold Hi-C lysis buffer (10 mM TRIS-HCl pH 8 (Sigma), 10 mM NaCl (Sigma), 0.2% Nonidet P-40 substitute (Sigma), 1 × complete EDTA-free protease inhibitor (Roche) for 30 min at 4 °C. The DNA was then digested using Dpn II (375 U DpnII (NEB, R0543 T/M/L) in 1× Dpn II buffer (NEB, B0543S) for 16 h at 37 °C with shaking at 800 rpm on a thermal block. Cells were centrifuged, the supernatant discarded, and fill-in mix added (38 µM biotin-14-dATP (Thermo Fisher Scientific, 19524016), 38 µM dCTP (Thermo Fisher Scientific, R0152), dGTP (Thermo Fisher Scientific, R0162), dTTP (Thermo Fisher Scientific, R0172), 50 U Klenow Polymerase (NEB, M0210L), 1 × NEB 2 buffer, B7002S). Cells were then incubated for 1 h at 37 °C with gentle rotation. Cells were then spun down, the supernatant discarded, ligation mix added (1 × T4 DNA ligase buffer (Thermo Fisher Scientific, B69), 0.1% Triton X-100 (Sigma, X100-100 ML), 100 µg mL-1 BSA (Sigma, A7030-10G), 50 U T4 DNA ligase (Thermo Fisher Scientific, EL0011) before incubation at room temperature for 4 h with gentle rotation. Cells were spun down, the supernatant discarded, and the pellet resuspended in 200 µL PBS. Genomic DNA (gDNA) was then purified using the DNeasy Blood and Tissue Kit (Qiagen, 69506) according to the manufacturer instructions for tissue culture cells. During gDNA purification, after addition of lysis buffer, RNaseA (Qiagen, 19101) and proteinase K (Qiagen, 19131), samples were incubated at 65 °C for 6 h to reverse crosslinks, before continuing with the rest of the protocol. The purified DNA was then sheared on a Covaris S2 instrument (duty cycle 10%, intensity 5.0, cycles/burst 200) for 25 s. The DNA was sheared in Covaris microTUBEs (Covaris, 520045). Size-selection was then performed to isolate fragments of the correct size using AMPure XP beads (Beckmann Coulter, A63881). A double size selection was performed according to manufacturer instructions. In brief 0.8 x sample volume of beads was first added to the beads, the supernatant was then transferred to a new tube and 0.12 x sample volume beads was then added. The final size-selected DNA was resuspended in dH_2_O. A biotin-pull down was then performed to select biotin-labelled DNA fragments using Dynabeads MyOne Streptavidin C1 beads (Thermo Fisher Scientific, 65001), resuspended in biotin binding buffer (5 mM TRIS-HCl pH 7.5 (Sigma), 0.5 mM EDTA (AppliChem, 1103.1000), 1M NaCl (Merck). Beads were incubated with the samples for 1 h at room temperature with gentle rotation before two washes with Tween wash buffer (5 mM TRIS-HCl, pH 7.5 (Sigma), 0.5 mM EDTA (Applichem, 1103.1000), 1M NaCl (Merck), 0.05% Tween20 (Sigma, P1379-250ML). A final wash with dH_2_O was then performed to remove residual detergent before the library preparation. The beads were then resuspended in dH_2_O, and library prep performed on the beads using the NEBNext Ultra II DNA library prep kit for Illumina (NEB, E7645L), according to the manufacturer instructions. The beads were then washed 4 x with Tween wash buffer. Elution of DNA from the beads was performed by incubation with 95% formamide (Sigma Aldrich, F9037-100ML), 10 mM EDTA (Applichem) at 65 °C for 2 min. Samples were then ethanol precipitated (80% EtOH (Sigma, 32221-1L / 2.5L), 100 mM sodium acetate pH 5.2 (Sigma Aldrich, 71180-1kg), incubation at -70 °C for > 16 h). 1 µL GlycoBlue (Thermo Fisher Scientific, AM9515) was also added during the precipitation to better visualise the pellet. Samples were spun down, the supernatant removed, washed once with 75% EtOH, spun down once more and then resuspended in dH_2_O. 4sT was then converted to methyl-cytosine using OsO_4_ / NH_4_Cl as described in (Mitter *et al*, 2020, 2022). EtOH precipitation was then performed once more to the converted samples as described above. Finally, qPCR was performed to amplify libraries using the NEBUltra Ultra II DNA library prep kit for Illumina (NEB, E7645L). The final libraries were purified using AMP Pure XP beads (Beckmann Coulter, A63881) at 0.55 x sample volume, according to the manufacturer instructions.

### Western blotting

For scsHi-C experiments, 5-10% of the harvested sample was set aside for Western blot analysis. 2 x Laemmli buffer (Bio-Rad, 1610737), 10 mM DTT (Roche, 10708984001) was added to the samples, before incubation at 95 °C for 10 min. For cells grown in 12-well or 24-well plates, Laemmli buffer was added directly to the well and the sample harvested, before incubation at 95 °C for 10 min. Before loading, samples were spun down once at 21000 g for 5 min. Samples were separated using Nu-Page 4-12% BisTris cassettes (Invitrogen, NP0326BOX) in 1x MES running buffer (Invitrogen, NP0002) according to the manufacturer instructions, apart from blots against NIPBL which were separated using 3-8% Tris-Acetate cassettes (Invitrogen, EA0375BOX) and 1 x Tris acetate running buffer (Invitrogen, LA0041) according to manufacturer instructions. Samples were then transferred to a 0.45 µm polyvinylidene (PVDF) membrane (activated using EtOH or MeOH) using a wet transfer (1 x transfer buffer, IMP/IMBA/GMI Molecular Biology Service/ Media Kitchen, 20% EtOH or MeOH, or 1 x NuPage™ transfer buffer, Thermo Fisher, NP0006-1, without methanol, for anti-NIPBL blots) for 1 h at room temperature. Membranes were blocked using 5% milk (w/v) (Maresi, “fixmilch” Instant milk powder), dissolved in PBS containing 0.05% Tween-20, for 30 min at room temperature. Primary and secondary antibodies were also diluted in 5% milk. Primary antibody incubations were carried out for > 16 h at 4°C, before 3 × 10 min washes with 5% milk 0.05% Tween-20 (Sigma, P1379-250ML). Secondary antibody incubations were carried out for 1 h at room temperature before 3 x washes with PBS containing 0.05% Tween-20. Proteins were detected as follows: SMC4 was detected using a monoclonal rabbit antibody (Abcam, ab229213, 1:2500). NIPBL was detected using a rat monoclonal antibody (Absea, 010702F01, 1:500). Sororin was detected using a custom rabbit antibody (1:500), kindly provided by J-M. Peters and previously published in (Mitter *et al*, 2020). WAPL was detected using a custom rabbit antibody (Peters lab internal ID A961, 1:1000), kindly provided by J-M. Peters and previously published in (Tedeschi *et al*, 2013) or a mouse monoclonal antibody (Santa-Cruz Biotechnology, sc-365189, 1:1000). GAPDH was detected using a rabbit polyclonal antibody (Abcam, ab9485, 1:2500). Vinculin was detected using a recombinant monoclonal rabbit antibody (Abcam, ab129002, 1:15000). The following horseradish peroxidase (HRP) conjugated antibodies were used against antibodies raised in the requisite organism: goat anti-rat HRP (Amersham, NA935, 1:2000), goat anti-mouse HRP (Bio-Rad, 1706516, 1:5000), goat anti-rabbit HRP (Bio-Rad, 1706515, 1:5000). Blots were incubated with Pierce ECL Plus Western Blotting Substrate (Thermo Fisher Scientific, 32132) or Clarity Max Western ECL Substrate (BioRad, 1705062) and then visualised on a Bio-Rad ChemiDoc imager.

### Flow cytometry – DNA content analysis and cell cycle stage verification

Flow cytometry was performed to assess the cell cycle stage of all scsHi-C samples generated during this study as described in (Mitter *et al*, 2020). All centrifugation steps were performed at 1100 g for 1 min. In brief, when scsHi-C samples were harvested, 10% of the cells were set aside for flow cytometry analysis. Cells were transferred to 15 mL Falcon tubes, washed once with PBS, plus/minus auxin/arresting compounds as appropriate, spun down and then fixed with 70% EtOH (Sigma Aldrich, 32221-2.5L) at 4 °C for > 30 min. Samples were spun down and then permeabilised on ice for 15 min with 0.25% Triton-X-100 (Sigma Aldrich, X100-100 ML). After spinning down, samples were incubated with a mouse monoclonal anti-phospho-H3Ser10 antibody (Millipore, 05-806, 0.25 µg per sample) diluted in PBS containing 1% BSA for 1 h at room temperature. Cells were spun down, washed once with PBS containing 1% BSA and then incubated with the secondary antibody, diluted as for the primary antibody (goat anti-mouse Alexa Fluor 488, Molecular Probes, A11001, 1:300) for 30 min at room temperature, protected from light. Cells were then stained with 200 µg mL-1 RNaseA (Qiagen, 19101), 50 µg mL-1 propidium iodide (Sigma Aldrich, 81845) for 30 min protected from light to determine the DNA content. Cells were analysed on either a FACS Canto II (BD Biosiences) or Penteon (Novacyte) instrument. Analysis was performed using FlowJo(v10). Cells were gated in the following way: Cells were gated by plotting FSC-A/SSC-A. Singlets were then gated by plotting FSC-A/ FSC-H, and then PI-A/PI-H. Cell cycle stage was then determined by a scatterplot of FITC (H3S10P signal) and PI (DNA content). An example of the gating strategy can be found in Supplementary Figure 2.

### Sequencing

All samples in this study were sequenced on an Illumina Novaseq 6000 instrument. Patterned SP flow cells were using with paired end sequencing (PE250 read mode).

### Hi-C data preprocessing

Preprocessing of scsHi-C samples was performed using a custom Nextflow pipeline, as described in (Mitter *et al*, 2020).

### Scaling plot and cis/trans sister contact ratio analyses

Scaling plots for cis and trans sister chromatid contacts were generated as described in (Mitter *et al*, 2020). In Extended Data Figure 6 scaling plot analysis was performed for all G2 conditions simultaneously before visualisation of each condition independently. As multiple samples were compared at the same time, downsampling was performed using the ngs package (https://github.com/gerlichlab/ngs) such that the total number of cis sister and trans sister contacts separated by more than 1 Kb was equal across different samples. The different prometaphase conditions were analysed in the same way. The cis/trans ratio for each condition was calculated by dividing the average cis contact probability at a given genomic distance by the average trans contact probability at the same genomic distance. Cis/trans ratios were then plotted against genomic distance. To calculate the genomic resolution score, the cis/trans ratio for each condition was first interpolated using the ‘CubicSpline’ operation from the scipy ‘interpolate’ module (Virtanen *et al*, 2020). The interpolated data was then smoothed using a Savitzy Golay filter. A threshold of 1.25 (slightly above the noise of the data) was then applied to the cis/trans ratio data. The genomic resolution score was calculated by determining the first genomic distance at which the cis/trans ratio threshold was reached.

### Sample number and statistical analyses

No statistical methods were used to predetermine sample size. To test significance, the robust, nonparametric Mann–Whitney U-test was used. The statistical tests used, and exact P values (wherever possible) are indicated in the figure legends. In some cases, the calculated P value was beyond the precision limit of the module used. In these cases, it is noted in the figure legend and an upper bound given. For each microscopy dataset generated in this study, at least two independent replicates were performed. The number of replicates and number of cells analysed for each condition is indicated in the requisite figure legend. For each condition analysed using scsHi-C in this study, at least two independent replicates were performed, and the number of replicates is indicated in the requisite figure legend. A detailed summary of the read statistics for each scsHi-C replicate can be found in Supplementary Tables S4-S10. All statistical tests were performed using scipy(Virtanen *et al*, 2020).

### Published datasets

All published datasets used in this study can be found in Supplementary Table S11.

## Data and code availability

All genomics datasets generated in this study will be uploaded to GEO. All data and code used in this manuscript will be made available on publication.

## Acknowledgements

The authors thank the IMBA/IMP/GMI BioOptics and Molecular Biology Service and the Vienna BioCenter Next Generation Sequencing facilities for technical support, I. Patten and M. Spicer for comments on the manuscript, and K. Nagasaka for advice on sister chromatid labelling and reagents for genome engineering. Research in the laboratory of D.W.G. has been supported by the Austrian Academy of Sciences, the Austrian Science Fund (FWF; Doktoratskolleg ‘Chromosome Dynamics’ DK W1238), the Vienna Science and Technology Fund (WWTF; projects LS17-003 and LS19-001), and the European Research Council (ERC) under the European Union’s Horizon 2020 research and innovation programme (grant agreement no. 101019039). Research in the laboratory of J.-M.P. has been supported by Boehringer Ingelheim, the Austrian Research Promotion Agency (Headquarter grant FFG-852936), the European Research Council under the European Union’s Horizon 2020 research and innovation programme GA No 693949, the Human Frontier Science Program (grant RGP0057/2018) and the Vienna Science and Technology Fund (grant LS19-029). J.-M.P. is also an adjunct professor at the Medical University of Vienna.

## Author contributions

P.B. and D.W.G. conceived the project. P.B. designed, performed, and analysed all experiments, except for NIPBL-AID scsHi-C, which was performed by Z.T., and wild type (G2) and Sororin-AID experiments, which were previously published in (Mitter *et al*, 2020). C.C.H.L and P.B. developed image analysis procedures. P.B., W.T., and C.B. generated endogenously tagged cell lines. D.W.G. and J-M.P. acquired funding. D.W.G. supervised the project. P.B. and D.W.G. wrote the manuscript.

---

**Figure EV1.**
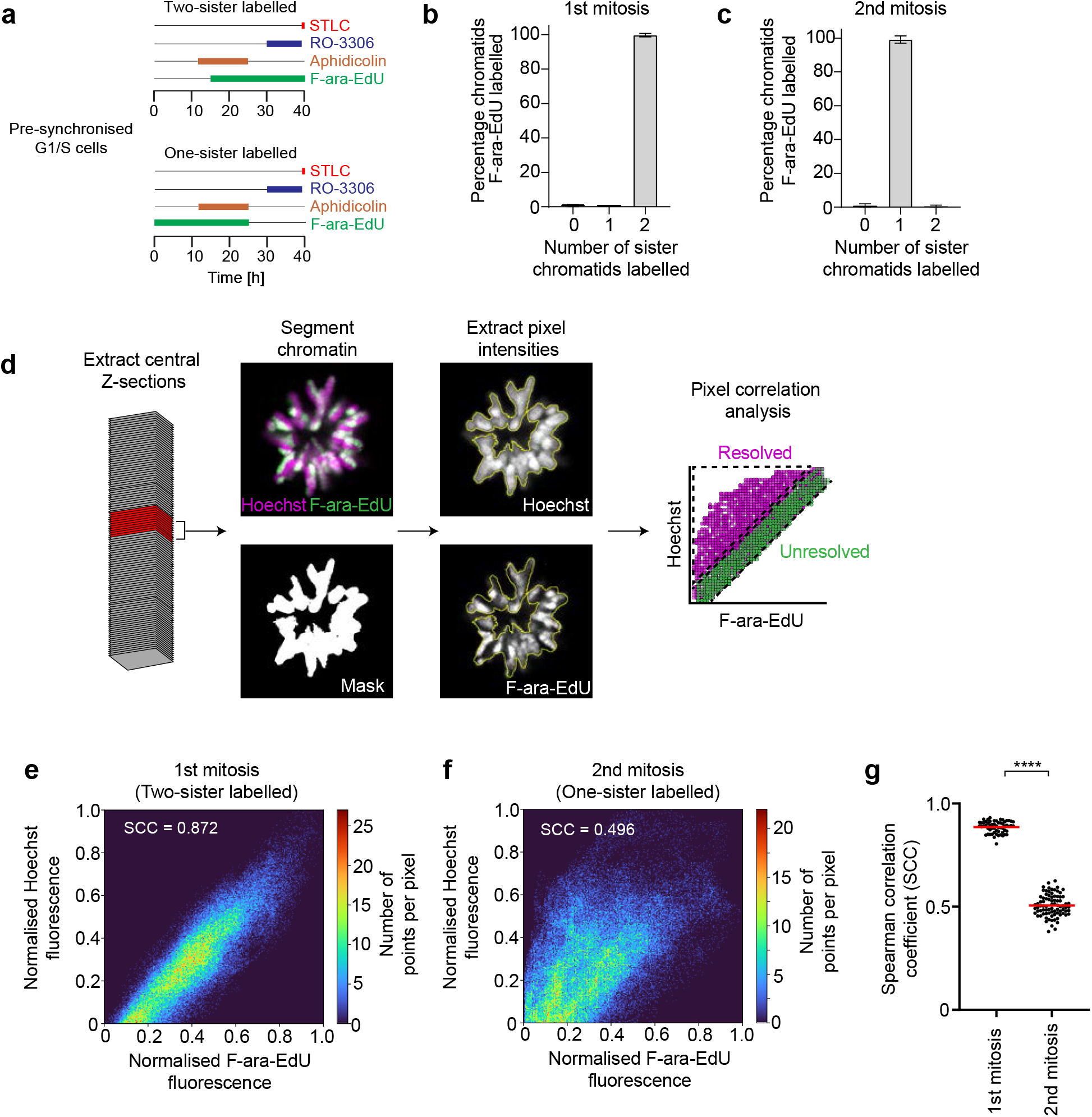
Validation of sister chromatid resolution assay. **a**, Schematic of experimental procedure for generation of two or one-sister labelled chromatids in wild type cells. Cells were treated with the requisite compounds as indicated. **b**, Bar plot indicating the percentage of chromosome segments with 0, 1 or 2 labelled sister chromatids for wild type prometaphase cells, fixed during the first mitosis after labelling with F-ara-EdU. n = 121 sister chromatid pairs from n = 15 cells analysed. Bar indicates mean; error bars indicate standard deviation. **c**, Bar plot indicating the percentage of chromosome segments with 0, 1 or 2 labelled sister chromatids for wild type prometaphase cells, fixed during the second mitosis after labelling with F-ara-EdU. n = 129 sister chromatid pairs from n = 27 cells analysed. Bar indicates mean; error bars indicate standard deviation. **d**, Schematic indicating the image analysis pipeline to calculate the sister chromatid separation score. Extract central Z-sections: The central slice of an input Z-stack is calculated, and the five slices above and below are chosen (11 slices in total). Segment chromatin, extract pixel intensities: The chromatin channel is segmented to generate a mask, which is applied to the Hoechst and F-ara-EdU channels. Pixel intensities within the mask are extracted. Pixel correlation analysis: The Spearman correlation coefficient (SCC) between Hoechst and F-ara-EdU pixel values in the mask is calculated first per slice, and then per cell. See Methods for details of the normalisation procedure to generate the separation score. Plot is an example scatterplot representation (real data not shown) of pixel value intensities for Hoechst and F-ara-EdU fluorescence in a prometaphase cell labelled on one sister chromatid. **e**, Scatterplot representation and SCC of pixel value intensities for Hoechst and F-ara-EdU fluorescence for a single slice of a wild type prometaphase cell, fixed during the 1^st^ mitosis after labelling with F-ara-EdU (two-sister labelled). Real data is shown. **f**, Scatterplot representation and SCC of pixel value intensities for Hoechst and F-ara-EdU fluorescence for a single slice of a wild type prometaphase cell, fixed during the 2^nd^ mitosis after labelling with F-ara-EdU (one-sister labelled). Real data is shown. **g**, Quantification of the Spearman correlation coefficient between Hoechst and F-ara-EdU for wild type prometaphase chromosomes fixed during the 1^st^ (two-sister labelled) or 2^nd^ (one-sister labelled) mitosis after labelling with F-ara-EdU, as indicated. Individual data points represent the mean SCC per cell, red bars indicate the mean. n = 59 cells from 8 experimental replicates (1^st^ mitosis), n = 83 cells from 14 experimental replicates (2^nd^ mitosis). Significance was tested using a two-tailed Mann Whitney U test; P = 3.91 × 10^−24^.

**Figure EV2.**
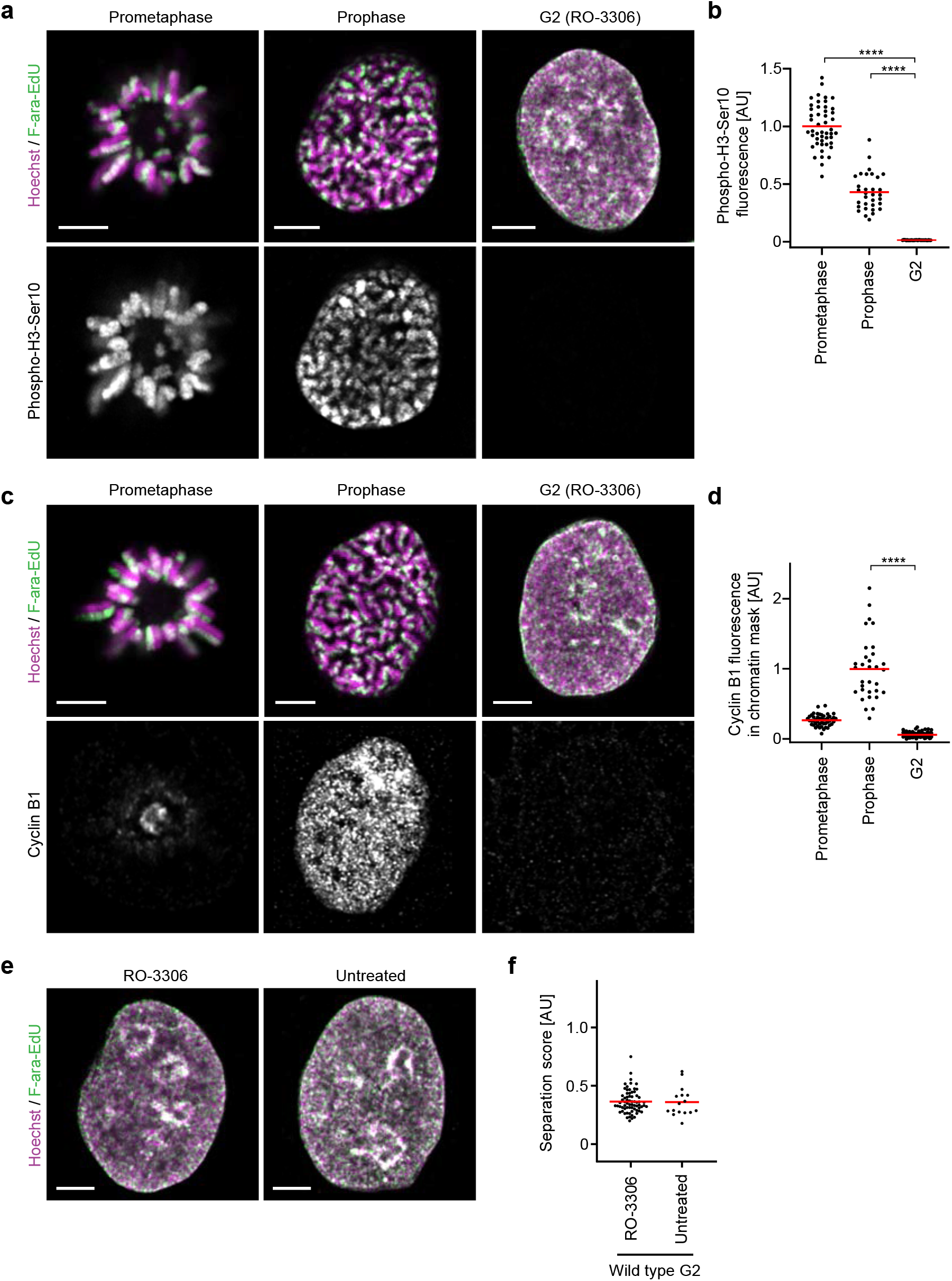
Validation of cell cycle stage for sister chromatid labelling. Cells were F-ara-EdU labelled on one sister chromatid and synchronised to different cell cycle stages as in Fig. 1 and 2 to determine the abundance and localisation of the mitotic histone modification phospho-H3-Ser10 and cyclin B1 by immunofluorescence. Prometaphase and prophase cells were treated with STLC, G2 cells were treated with RO-3306. **a**, Representative images for one-sister labelled sister chromatids in cells stained with an anti-phospho-H3-Ser10 antibody from wild type prometaphase, prophase and G2 cells as indicated. **b**, Quantification of mean phospho-H3-Ser10 fluorescence within the chromatin mask for central Z-slices for the conditions shown in **a**. n = 48 cells (prometaphase) from 4 experimental replicates, n = 32 cells (prophase) from 4 experimental replicates, n = 31 cells (G2) from 3 experimental replicates were analysed. Significance was tested using a two-tailed Mann Whitney U test; P = 9.61 × 10^−12^ (prophase), P = 8.33 × 10^−14^ (prometaphase). **c**, Representative images for one-sister labelled sister chromatids in cells stained with an anti-cyclin B1 antibody, from wild type prometaphase, prophase and G2 arrested cells as indicated. **d**, Quantification of mean cyclin B1 fluorescence within the chromatin mask for central Z-slices for the conditions shown in **c**. Calculation of mean fluorescence was performed as described in **b**. n = 48 cells (prometaphase) from 4 experimental replicates, n = 32 cells (prophase) from 4 experimental replicates, n = 69 cells (G2) from 8 experimental replicates. Significance was tested using a two-tailed Mann Whitney U test; P = 7.95 × 10^−16^ (prophase). **e**, Representative images for one-sister labelled sister chromatids from G2 cells, treated with RO-3306 or untreated, as indicated. **f**, Quantification of sister chromatid separation for one-sister labelled wild type G2 cells, as shown in **e**. RO-3306 treated (n = 69 cells, from 8 experimental replicates) and untreated (n = 16 cells from 2 experimental replicates) cells were analysed. Quantification for RO-3306 treated cells is the same as in Fig. 1e (wild type) to allow side by side comparison with untreated cells. A different example RO-3306 treated cell is shown. All images shown are single Z-slices from Z-stack images. Scale bars: 5 µm.

**Figure EV3.**
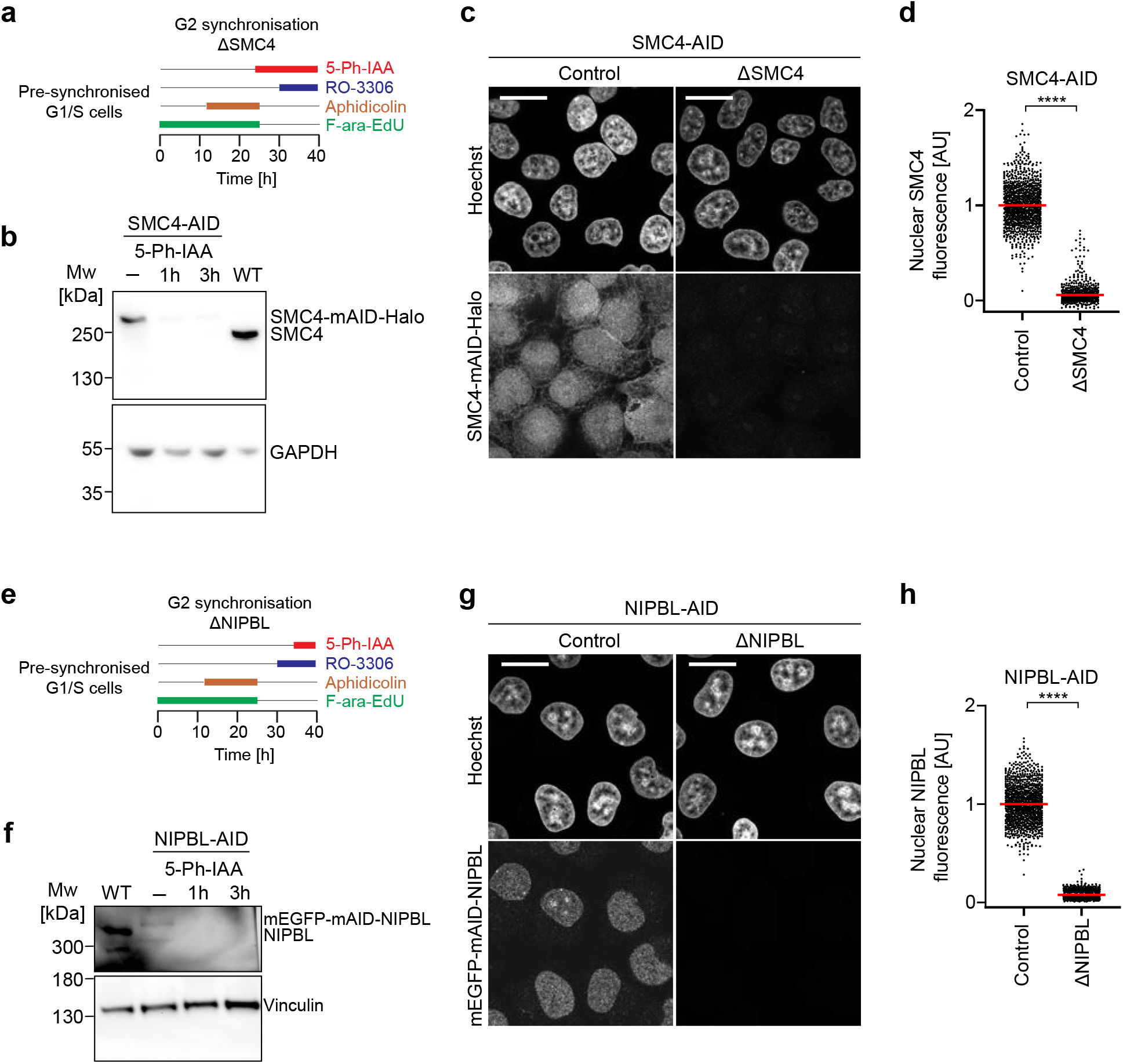
Validation of protein depletion efficiency in SMC4-AID and NIPBL-AID cells. **a**, Schematic of experimental procedure for generation of one-sister labelled chromatids in ΔSMC4 G2 cells. Cells were treated with the requisite compounds as indicated; the end point indicates the time of fixation. Prometaphase samples were generated by washing out RO-3306 and releasing into medium containing STLC. **b**, Immunoblot analysis of SMC4 in wild type (WT) cells, untreated SMC4-AID cells and SMC4-AID cells treated for either 1 h or 3 h with 5-Ph-IAA. Representative example of n = 3 experiments. For source gel data, see Supplementary Figure 1a. **c**, Immunofluorescence analysis of HeLa cells homozygously tagged for SMC4-mAID-Halo and stably expressing OsTIR1^F74G^. Cells were incubated for 3 h with (ΔSMC4) or without (Control) 1 µM 5-Ph-IAA before subsequently staining of SMC4 with TMR-HaloTag ligand. DNA was stained with Hoechst 33342. **d**, Quantification of mean nuclear SMC4 fluorescence per cell, as shown in **c**. Wild type cells were stained with Halo-TMR and the mean Halo-TMR fluorescence within the segmented nuclei then calculated. Normalisation was performed relative to the mean nuclear Halo-TMR fluorescence of wild type cells (0 value) and control SMC4-AID cells (1 value). For each condition two experimental replicates were performed. n = 1159 cells analysed for control SMC4-AID cells, n = 1000 cells analysed for ΔSMC4 cells. Significance was tested using a two-tailed Mann Whitney U test; P < 10^−324^ (precision limit of floating-point arithmetic). **e**, Schematic of experimental procedure for generation of one-sister labelled chromatids in ΔNIPBL G2 cells. Cells were treated with the requisite compounds as indicated; the end point indicates the time of fixation. Prometaphase samples were generated by washing out RO-3306 and releasing into medium containing STLC. **f**, Immunoblot analysis of NIPBL in wild type (WT) cells, untreated NIPBL-AID cells and NIPBL-AID cells treated for either 1 h or 3 h with 5-Ph-IAA. Representative example of n = 2 experiments. **g**, Immunofluorescence analysis of HeLa cells homozygously tagged for mEGFP-mAID-NIPBL and stably expressing OsTIR1^F74G^. Cells were incubated for 2 h with (ΔNIPBL) or without (Control) 1 µM 5-Ph-IAA before subsequently fixing and staining for mEGFP with an anti-GFP nanobody. DNA was stained with Hoechst 33342. **h**, Quantification of mean nuclear mEGFP-mAID-NIPBL fluorescence per cell, as shown in **g**. For each condition three experimental replicates were performed. n = 1205 cells analysed for control NIPBL-AID cells, n = 1223 cells analysed for ΔNIPBL cells. Significance was tested using a two-tailed Mann Whitney U test; P < 10^−324^ (precision limit of floating-point arithmetic). For source gel data, see Supplementary Figure 1b-c. All microscopy images are single Z-sections. Scale bars: 20 µm.

**Figure EV4.**
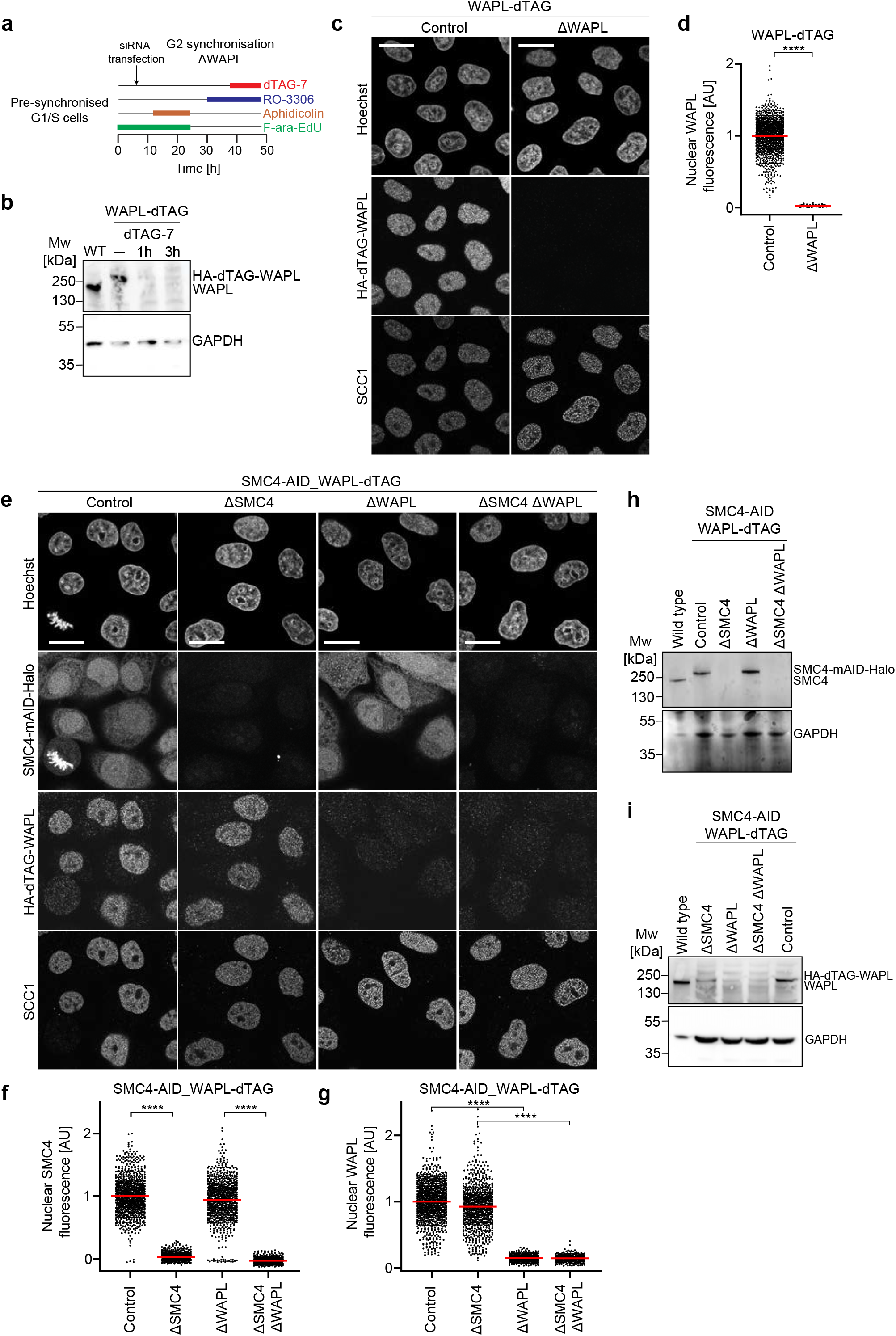
Validation of protein depletion efficiency in WAPL-dTAG and SMC4-AID/WAPL-dTAG cells. **a**, Schematic of experimental procedure for generation of one-sister labelled chromatids in ΔWAPL G2 cells. Cells were treated with the requisite compounds as indicated; the end point indicates the time of fixation. For conditions where WAPL was not depleted, dTAG-7 was not added to the cells. **b**, Immunoblot analysis of WAPL in wild type (WT) cells, untreated WAPL-dTAG cells and WAPL-dTAG cells treated for either 1 h or 3 h with dTAG-7. Representative example of n = 2 experiments. For source gel data, see Supplementary Figure 1d. **c**, Immunofluorescence analysis of HeLa cells homozygously tagged for HA-dTAG-WAPL. Cells were incubated for 3 h with (ΔWAPL) or without (Control) 1 µM dTAG-7 before subsequently fixing and staining for HA-WAPL and SCC1, with anti-HA and anti-SCC1 antibodies respectively. DNA was stained with Hoechst 33342. **d**, Quantification of mean nuclear HA-WAPL fluorescence per cell, as shown in **c**. For each condition two experimental replicates were performed. n = 1466 cells analysed for control WAPL-dTAG cells, n = 1372 cells analysed for ΔWAPL cells. Significance was tested using a two-tailed Mann Whitney U test; P < 10^−324^ (precision limit of floating-point arithmetic). **e**, Immunofluorescence analysis of HeLa cells homozygously tagged for SMC4-mAID-Halo and HA-dTAG-WAPL, and stably expressing OsTIR1^F74G^. Cells were incubated with 5-Ph-IAA or dTAG-7 in the following combinations: untreated (Control), 3 h 5-Ph-IAA (ΔSMC4), 3 h dTAG-7 (ΔWAPL), 3 h 5-Ph-IAA + dTAG-7 (ΔSMC4 ΔWAPL). SMC4 was stained with TMR-HaloTag ligand and HA-WAPL and SCC1 were visualised using antibodies as in **a**. DNA was stained with Hoechst 33342. **f**, Quantification of mean nuclear SMC4 fluorescence per cell, as shown in **e**. Wild type cells were stained with Halo-TMR and the mean Halo-TMR fluorescence within the segmented nuclei then calculated. Normalisation was performed relative to the mean nuclear Halo-TMR fluorescence of wild type cells (0 value), and control SMC4-AID_WAPL-dTAG cells (1 value). For each condition two experimental replicates were performed. n = 1007 cells analysed for control SMC4-AID_WAPL-dTAG cells, n = 848 cells analysed for ΔSMC4 cells, n = 867 cells analysed for ΔWAPL cells, n = 717 cells analysed for ΔSMC4 ΔWAPL cells. Significance was tested using a two-tailed Mann Whitney U test; P = 1.07 × 10^−297^ (ΔSMC4), P = 3.81 × 10^−244^ (ΔSMC4 ΔWAPL). **g**, Quantification of mean nuclear HA-WAPL fluorescence per cell, as shown in **e**. Sample numbers as in **f**. Significance was tested using a two-tailed Mann Whitney U test; P = = 6.36 × 10^−305^ (ΔWAPL), P = 5.15 × 10^−253^ (ΔSMC4 ΔWAPL). **h**, Immunoblot analysis of SMC4 in wild type and SMC4-AID_WAPL-dTAG cells treated as follows: untreated (Control), 3 h 5-Ph-IAA (ΔSMC4), 3 h dTAG-7 (ΔWAPL), 3 h 5-Ph-IAA + 3 h dTAG-7 (ΔSMC4 ΔWAPL). Representative example of n = 2 experiments. For source gel data, see Supplementary Figure 1e-f. **i**, Immunoblot analysis of WAPL in wild type and SMC4-AID_WAPL-dTAG cells treated as follows: untreated (Control), 3 h 5-Ph-IAA (ΔSMC4), 3 h dTAG-7 (ΔWAPL), 3 h 5-Ph-IAA + 3 h dTAG-7 (ΔSMC4 ΔWAPL). Representative example of n = 2 experiments. For source gel data, see Supplementary Figure 1g. All microscopy images are single Z-sections. Scale bars: 20 µm.

**Figure EV5.**
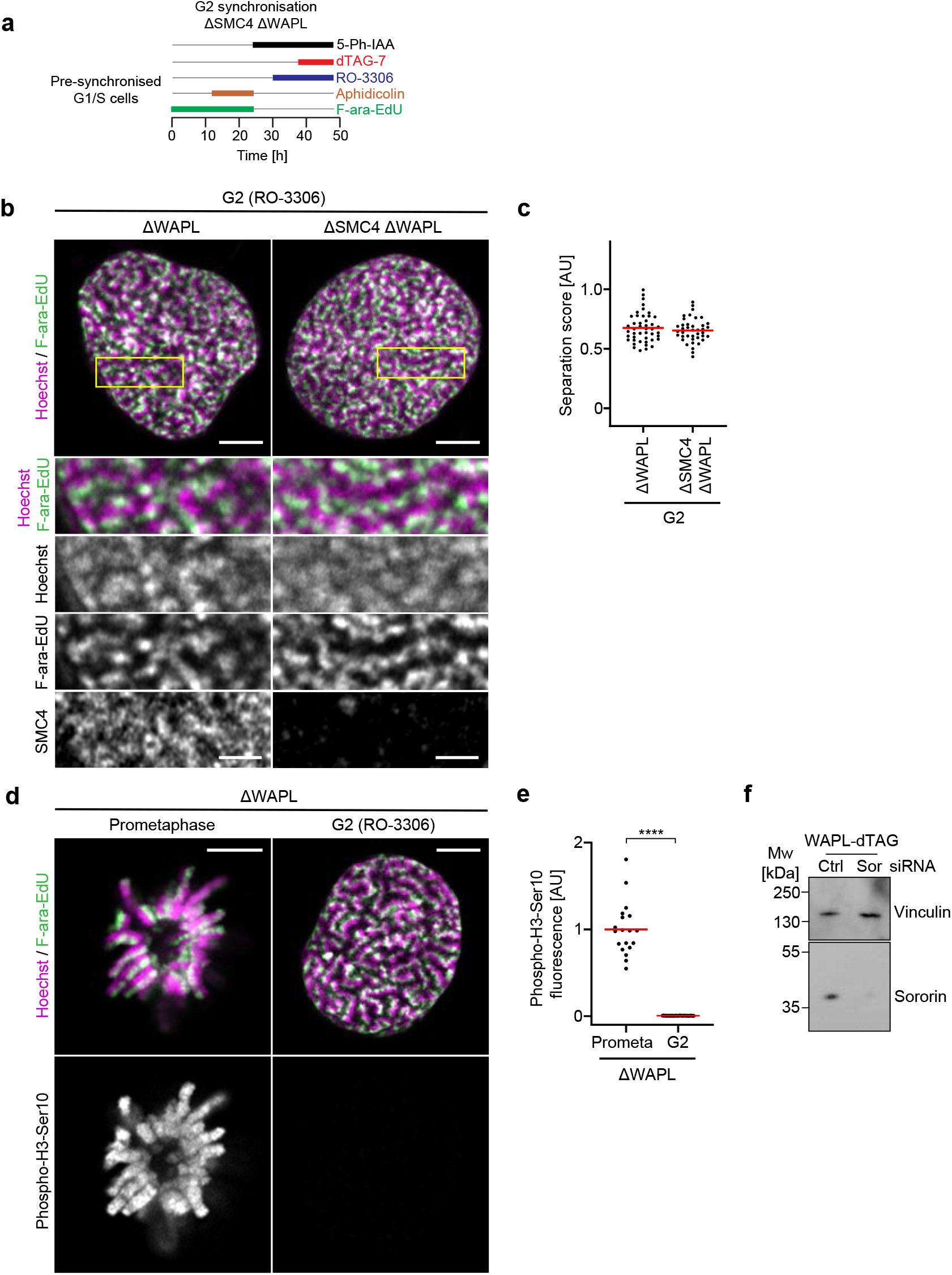
Condensin is not required to resolve sister DNAs in WAPL depleted G2 cells. **a**, Schematic of experimental procedure for generation of one-sister labelled chromatids in ΔSMC4 ΔWAPL G2 cells. Cells were treated with the requisite compounds as indicated; the end point indicates the time of fixation. **b**, Representative images of one-sister labelled sister chromatids from ΔWAPL or ΔSMC4 ΔWAPL G2 cells, as indicated. Cells were labelled and fixed as in Fig. 2b. SMC4 was depleted in G1 through the addition of 5-Ph-IAA 1 h before the final S phase release. SMC4 was visualised by staining with TMR-HaloTag ligand. Wild type cells were stained with Halo-TMR to determine the fluorescence background for background subtraction. The SMC4 channel is displayed after background subtraction. **c**, Separation score quantification for one-sister labelled ΔWAPL G2 (n = 45 cells from 6 experimental replicates) and ΔSMC4 ΔWAPL G2 cells (n = 40 cells from 5 experimental replicates). Dots represent individual cells; red bars indicate the mean. **d**, Validation of cell cycle stage in ΔWAPL cells by phospho-H3-Ser10 immunofluorescence in cells arrested in prometaphase (prometa) by STLC and cells arrested in G2 by RO-3306. **e**, Quantification of mean phospho-H3-Ser10 fluorescence for central Z-stack slices for the conditions shown in **d**, as in Extended Data Fig. 2b. n = 19 cells from 3 experimental replicates (ΔWAPL prometaphase), n = 33 cells from 3 experimental replicates (ΔWAPL G2). Significance was tested using a two-tailed Mann Whitney U test; P = 2.72 × 10^−9^. **f**, Immunoblot analysis of Sororin in WAPL-dTAG cells treated with either Control (Ctrl) or Sororin (Sor) siRNAs as indicated, as in Fig. 2e. Cells were harvested 40 h after transfection. Representative example of n = 2 experiments. For source gel data, see Supplementary Figure 1h. All microscopy images are single Z-sections. Yellow boxes indicate inset regions. Scale bars large panels: 5 µm, insets: 2 µm.

**Figure EV6.**
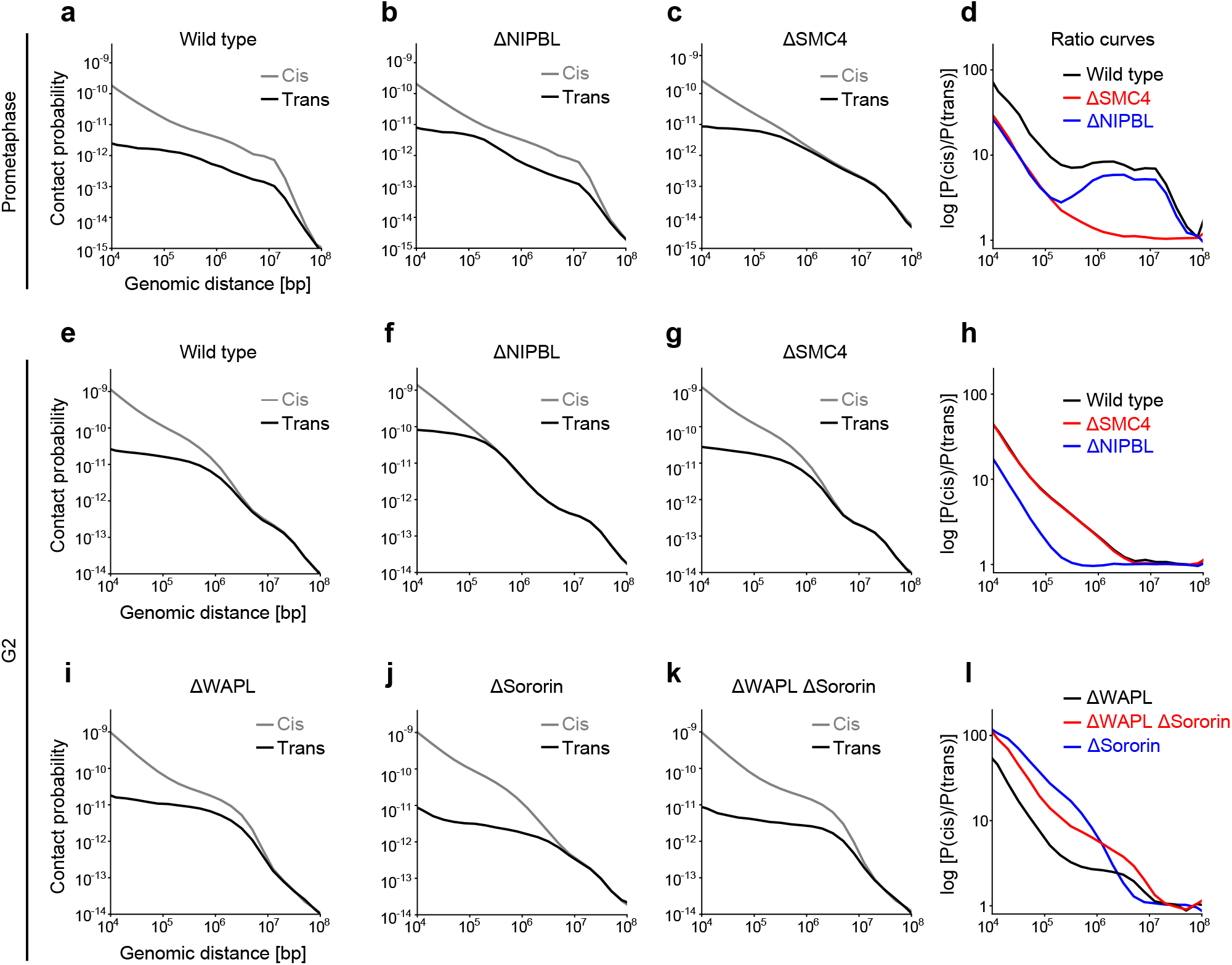
scsHi-C contact probability curves. Sister-chromatid-sensitive Hi-C experiments were analysed by calculating the average contact probability for cis sister and trans sister contacts over variable genomic intervals. **a**, Cis sister and trans sister contact probability curves for wild type prometaphase cells (merge of 2 biologically independent experiments). **b**, Cis sister and trans sister contact probability curves for ΔNIPBL prometaphase cells (merge of 3 biologically independent experiments). **c**, Cis sister and trans sister contact probability curves for ΔSMC4 prometaphase cells (merge of 2 biologically independent experiments). **d**, Ratio curves plotting cis sister contact probability / trans sister contact probability against genomic distance for the conditions in **a-c**. Merged curves of the individual replicates shown. **e**, Cis sister and trans sister contact probability curves for wild type G2 cells (merge of 11 biologically independent experiments). **f**, Cis sister and trans sister contact probability curves for ΔNIPBL G2 cells (merge of 10 biologically independent experiments). **g**, Cis sister and trans sister contact probability curves for ΔSMC4 G2 cells (merge of 4 biologically independent experiments). **h**, Ratio curves plotting cis sister contact probability / trans sister contact probability against genomic distance for the conditions in **e-g**. Merged curves of the individual replicates shown. **i**, Cis sister and trans sister contact probability curves for ΔWAPL G2 cells (merge of 6 biologically independent experiments). **j**, Cis sister and trans sister contact probability curves for ΔSororin G2 cells (merge of 3 biologically independent experiments). **k**, Cis sister and trans sister contact probability curves for ΔWAPL ΔSororin G2 cells (merge of 4 biologically independent experiments). **l**, Ratio curves plotting cis sister contact probability / trans contact probability against genomic distance for the conditions in **i-k**. Merged curves of the individual replicates shown.

**Figure EV7.**
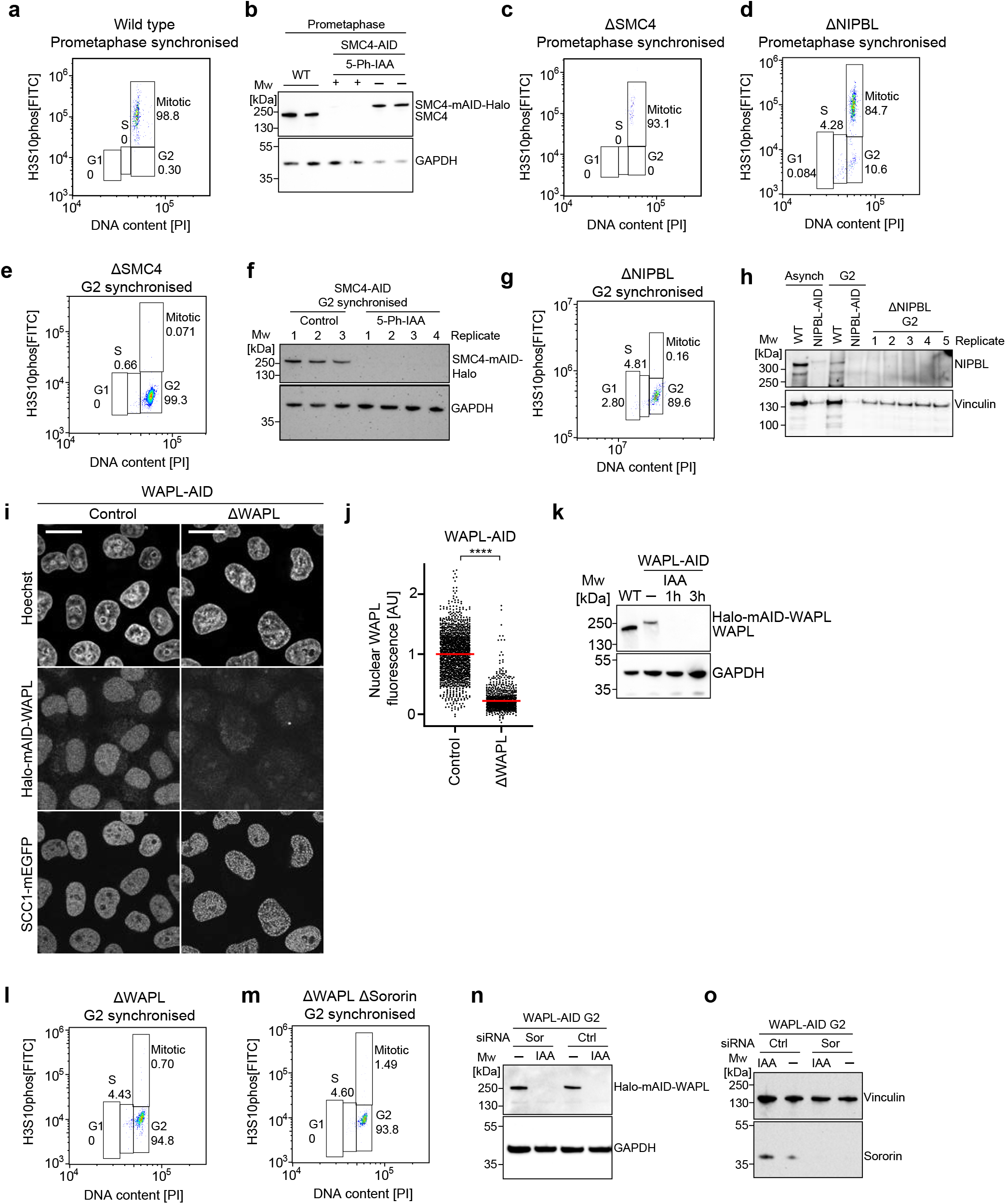
Validation of cell cycle stage and protein depletion efficiency for scsHi-C. a, c, d, e, g, l, m, Cell cycle analysis by flow cytometry. Cells were stained with propidium iodide to determine DNA content and an antibody against phospho-H3-Ser10 as a marker for mitotic cells. The gates indicate the cell cycle stage, and the numbers indicate the percentage of cells measured for a given cell cycle stage. **a**, Representative flow cytometry plot for one of the wild type prometaphase samples (n = 2 replicates). **b**, Immunoblot analysis of SMC4 in wild type (WT) and SMC4-AID prometaphase cells harvested for scsHi-C analysis, plus or minus 5-Ph-IAA as indicated. For source gel data, see Supplementary Figure 1i. **c**, Representative flow cytometry plot for one of the ΔSMC4 prometaphase samples (n = 2 replicates). **d**, Representative flow cytometry plot for one of the ΔNIPBL prometaphase samples (n = 3 replicates). **e**, Representative flow cytometry plot for one of the ΔSMC4 G2 samples (n = 4 replicates). **f**, Immunoblot analysis of G2 synchronised control SMC4-AID and ΔSMC4 cells harvested for scsHi-C analysis, plus or minus 5-Ph-IAA as indicated. For source gel data, see Supplementary Figure 1j. **g**, Representative flow cytometry plot for one of the ΔNIPBL G2 samples (n = 10 replicates). **h**, Immunoblot analysis of NIPBL in wild type (WT) and G2 synchronised NIPBL-AID cells harvested for downstream scsHi-C analysis. For source gel data, see Supplementary Figure 1k. **i**, Immunofluorescence analysis of HeLa cells homozygously tagged for Halo-mAID-WAPL and stably expressing OsTIR1. Cells were incubated for 2 h with (ΔWAPL) or without (Control) 500 µM auxin before subsequently staining of WAPL with TMR-HaloTag ligand. DNA was stained with Hoechst 33342. Images show single Z-sections. **j**, Quantification of mean nuclear WAPL fluorescence per cell, as shown in **i**. Wild type cells were stained with Halo-TMR and the mean Halo-TMR fluorescence within the segmented nuclei then calculated. Normalisation was performed relative to the mean nuclear Halo-TMR fluorescence of wild type cells (0 value) and control WAPL-AID cells (1 value). For each condition three experimental replicates were performed. n = 1537 cells analysed for control WAPL-AID cells, n = 1199 cells analysed for ΔWAPL cells. Significance was tested using a two-tailed Mann Whitney U test; P = < 10^−324^ (precision limit of floating-point arithmetic). **k**, Immunoblot analysis of WAPL in wild type cells, untreated WAPL-AID cells and WAPL-AID cells treated for either 1 h or 3 h with auxin (IAA). Representative example of n = 3 experiments. For source gel data, see Supplementary Figure 1l. **l**, Representative flow cytometry plot for one of the ΔWAPL G2 samples (n = 6 replicates). **m**, Representative flow cytometry plot for one of the ΔWAPL ΔSororin G2 samples (n = 4 replicates). **n**, Immunoblot analysis of WAPL in WAPL-AID cells harvested for scsHi-C analysis, treated with Control (Ctrl) or Sororin (Sor) siRNAs, plus or minus auxin (IAA), as indicated. For source gel data, see Supplementary Figure 1m. **o**, Immunoblot analysis of Sororin in WAPL-AID cells harvested for scsHi-C analysis, treated with Control (Ctrl) or Sororin (Sor) siRNAs, plus or minus auxin (IAA) as indicated. For source gel data, see Supplementary Figure 1n.

**Supplementary Figure 1.**
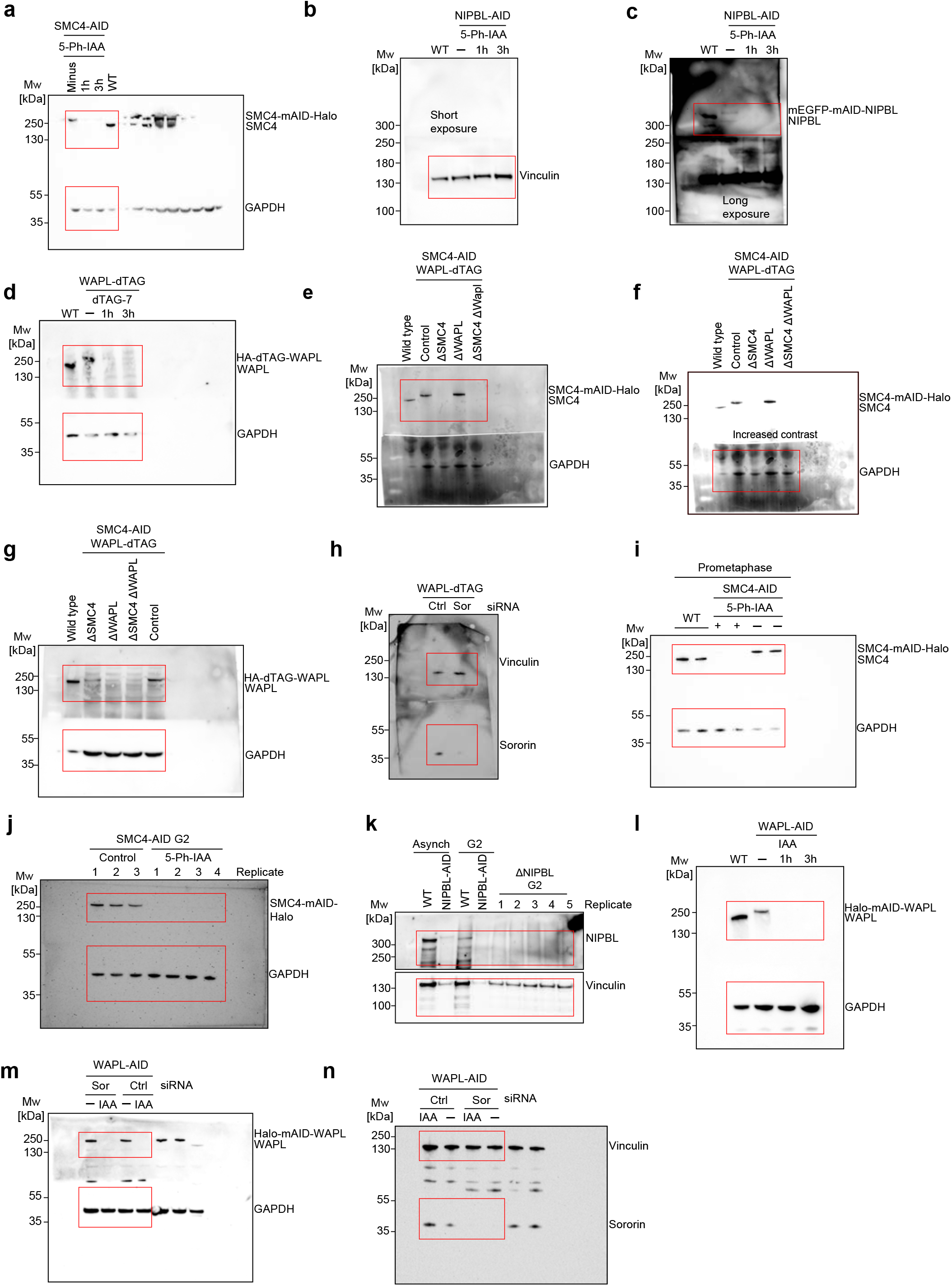
Uncropped Western blot images. Loading controls were run on the same gel as the target protein, and unless otherwise specified, the membrane was cut at 70 KDa after transfer to allow detection of different proteins. In all cases, the cropped images used in the requisite figure panels are indicated with red boxes. **a**, Characterisation of SMC4-AID cells. Uncropped blot of wild type (WT) and SMC4-AID cells (untreated, 1 h 5-Ph-IAA, 3 h 5-Ph-IAA), as indicated, blotting against SMC4 and GAPDH. Cropped images shown in Extended Data Figure 3c. **b**, Characterisation of NIPBL-AID cells. Uncropped blot of wild type (WT) and NIPBL-AID cells (untreated, 1 h 5-Ph-IAA, 3 h 5-Ph-IAA, as indicated), short exposure time, showing the loading control (Vinculin). Cropped area shown in Extended Data Figure 3f. Loading controls were run on the same gel and the membrane cut after transfer at 250 KDa to stain NIPBL and Vinculin with different antibodies **c**, Uncropped blot as in **b**, long exposure time, blotting against NIPBL. **d**, Characterisation of WAPL-dTAG cells. Uncropped blot of wild type (WT) and WAPL-dTAG cells (untreated, 1 h dTAG-7, 3 h dTAG-7, as indicated). Cropped area shown in Extended Data Figure 4c, blotting against WAPL and GAPDH. **e**, Characterisation of SMC4-AID_WAPL-dTAG cells. Uncropped blot of wild type and SMC4-AID_WAPL-dTAG cells (untreated, 3 h 5-Ph-IAA, 3 h dTAG-7, 3 h 5-Ph-IAA + dTAG-7, as indicated), showing blotting against SMC4. Cropped area shown in Extended Data Figure 4g, blotting against SMC4 and GAPDH. **f**, Uncropped blot as in **e**, increased contrast, showing blotting against the loading control (GAPDH). **g**, Characterisation of SMC4-AID_WAPL-dTAG cells. Same conditions as in **e-f**, blotting against WAPL and GAPDH. Cropped areas shown in Extended Data Figure 4h, blotting against WAPL and GAPDH. **h**, Uncropped blot of WAPL-dTAG cells treated with Control (Ctrl) or Sororin (Sor) siRNAs, as indicated. Cropped areas shown in Extended Data Figure 5e, blotting against Sororin and Vinculin. **i**, Uncropped blot of prometaphase synchronised wild type (WT) and SMC4-AID cells harvested for scsHi-C analysis, plus or minus 5-Ph-IAA as indicated. Cropped areas shown in Extended Data Figure 7b, blotting against SMC4 and GAPDH. **j**, Uncropped blot of G2 synchronised control SMC4-AID and ΔSMC4 cells harvested for scsHi-C analysis, plus or minus 5-Ph-IAA as indicated. Cropped areas shown in Extended Data Figure 7f, blotting against SMC4 and GAPDH. **k**, Uncropped blot of wild type (WT) and G2 synchronised NIPBL-AID cells harvested for scsHi-C analysis. Cropped areas shown in Extended Data Figure 7h, blotting against NIPBL and Vinculin. **l**, Characterisation of WAPL-AID cell line. Uncropped blot of wild type (WT) and WAPL-AID cells (untreated, 1 h auxin (IAA), 3 h auxin (IAA), as indicated). Cropped area shown in Extended Data Figure 7k, blotting against WAPL and GAPDH. **m**, Uncropped blot of G2 synchronised WAPL-AID cells harvested for scsHi-C analysis, treated with Control (Ctrl) or Sororin (Sor) siRNAs, plus or minus auxin (IAA), as indicated. Cropped areas shown in Extended Data Figure 7n, blotting against WAPL and GAPDH. **N**, Uncropped blot of G2 synchronised WAPL-AID cells harvested for scsHi-C analysis, treated with Control (Ctrl) or Sororin (Sor) siRNAs, plus or minus auxin (IAA), as indicated. Cropped areas shown in Extended Data Figure 7o, blotting against Sororin and Vinculin.

**Supplementary Figure 2.**
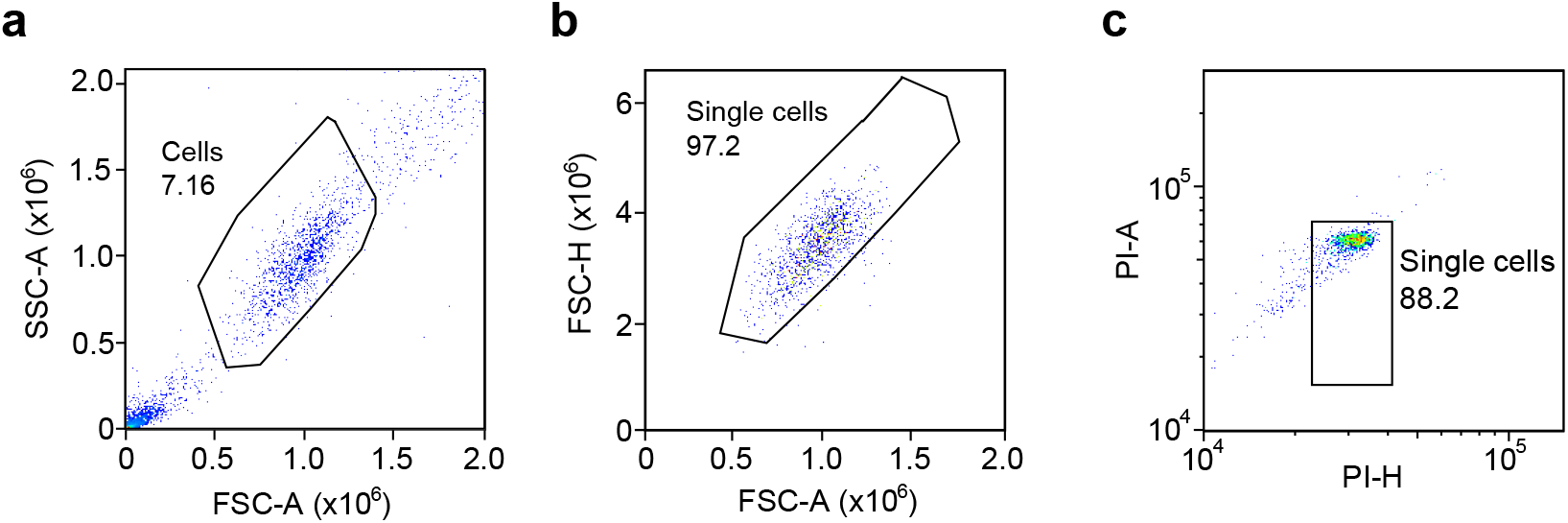
Flow cytometry gating strategy. For all panels, numbers indicate the percentage of cells inside the gate. Cells were stained with propidium iodide to determine DNA content and an antibody against phospho-H3-Ser10 as a marker for mitotic cells. **a**, FSC-A vs SSC-A gate to gate cells and remove debris, shown for one of the G2 synchronised ΔWAPL samples. **b**, FSC-A vs FSC-H gate to identify single cells. **c**, PI-H vs PI-A, second gate to identify single cells. Cells in this gate were subsequently used to analyse cell cycle stage in Extended Data Fig.7l. The same gating strategy was used for Extended Data Fig. 7a, c, d, e, g, m.

**Supplementary Table S1.** Flow cytometry statistics for G2 scsHi-C samples.

| Condition | Replicate | #G1 | #S | #G2 | #M | % G1 | % S | % G2 | %M |
| --- | --- | --- | --- | --- | --- | --- | --- | --- | --- |
| ΔWAPL G2 | Replicate 1 | 87 | 57 | 1151 | 9 | 6.64 | 4.35 | 87.9 | 0.69 |
| ΔWAPL G2 | Replicate 2 | 4 | 5 | 260 | 1 | 1.48 | 1.85 | 95.9 | 0.37 |
| ΔWAPL G2 | Replicate 3 | 0 | 23 | 329 | 1 | 0 | 6.53 | 93.5 | 0.28 |
| ΔWAPL G2 | Replicate 4 | 0 | 51 | 1095 | 8 | 0 | 4.42 | 94.8 | 0.69 |
| ΔWAPL G2 | Replicate 5 | 1 | 107 | 1632 | 12 | 0 | 6.1 | 93 | 0.68 |
| ΔWAPL G2 | Replicate 6 | 0 | 53 | 613 | 9 | 0 | 7.83 | 90.5 | 1.33 |
| ΔWAPL ΔSororin G2 | Replicate 1 | 5 | 7 | 430 | 1 | 1.13 | 1.58 | 97.1 | 0.23 |
| ΔWAPL ΔSororin G2 | Replicate 2 | 0 | 74 | 541 | 4 | 0 | 11.9 | 87.1 | 0.64 |
| ΔWAPL ΔSororin G2 | Replicate 3 | 0 | 90 | 992 | 2 | 0 | 8.29 | 91.3 | 0.18 |
| ΔWAPL ΔSororin G2 | Replicate 4 | 0 | 91 | 594 | 8 | 0 | 13.1 | 85.3 | 1.15 |
| ΔSMC4 G2 | Replicate 1 | 0 | 14 | 1264 | 0 | 0 | 1.1 | 99.1 | 0 |
| ΔSMC4 G2 | Replicate 2 | 0 | 17 | 1317 | 0 | 0 | 1.28 | 98.9 | 0 |
| ΔSMC4 G2 | Replicate 3 | 0 | 7 | 1009 | 0 | 0 | 0.69 | 99.3 | 0 |
| ΔSMC4 G2 | Replicate 4 | 0 | 28 | 4195 | 3 | 0 | 0.66 | 99.3 | 0.071 |
| ΔNIPBL G2 | Replicate 1 | 18 | 31 | 577 | 1 | 2.8 | 4.81 | 89.6 | 0.16 |
| ΔNIPBL G2 | Replicate 2 | 17 | 56 | 661 | 2 | 2.25 | 7.41 | 87.4 | 0.26 |
| ΔNIPBL G2 | Replicate 3 | 18 | 50 | 518 | 1 | 3.01 | 8.35 | 86.5 | 0.17 |
| ΔNIPBL G2 | Replicate 4 | 20 | 67 | 665 | 1 | 2.6 | 8.7 | 86.4 | 0.13 |
| ΔNIPBL G2 | Replicate 5 | 18 | 65 | 816 | 2 | 1.97 | 7.12 | 89.4 | 0.22 |
| ΔNIPBL G2 | Replicate 6 | 52 | 112 | 1172 | 1 | 3.81 | 8.2 | 85.8 | 0.073 |
| ΔNIPBL G2 | Replicate 7 | 60 | 148 | 1466 | 0 | 3.5 | 8.64 | 85.6 | 0 |
| ΔNIPBL G2 | Replicate 8 | 43 | 83 | 874 | 0 | 4.2 | 8.1 | 85.3 | 0 |
| ΔNIPBL G2 | Replicate 9 | 36 | 87 | 889 | 0 | 3.44 | 8.32 | 85 | 0 |
| ΔNIPBL G2 | Replicate 10 | 96 | 136 | 1533 | 0 | 5.33 | 7.55 | 85.1 | 0 |

**Supplementary Table S2.** Flow cytometry statistics for prometaphase scsHi-C samples.

| Condition | Replicate | #G1 | #S | #G2 | #M | % G1 | % S | % G2 | %M |
| --- | --- | --- | --- | --- | --- | --- | --- | --- | --- |
| WT prometaphase | Replicate 1 | 0 | 4 | 5 | 150 | 0 | 2.44 | 3.05 | 91.05 |
| WT prometaphase | Replicate 2 | 0 | 0 | 2 | 666 | 0 | 0 | 0.3 | 98.8 |
| ΔSMC4 prometaphase | Replicate 1 | 0 | 0 | 2 | 115 | 0 | 0 | 1.64 | 94.3 |
| ΔSMC4 prometaphase | Replicate 2 | 0 | 0 | 0 | 147 | 0 | 0 | 0 | 93.1 |
| ΔNIPBL prometaphase | Replicate 1 | 1 | 52 | 126 | 1010 | 0.084 | 4.36 | 10.6 | 84.7 |
| ΔNIPBL prometaphase | Replicate 2 | 0 | 30 | 87 | 556 | 0 | 4.44 | 12.9 | 82.2 |
| ΔNIPBL prometaphase | Replicate 3 | 0 | 52 | 82 | 707 | 0 | 6.09 | 10.3 | 82.8 |

**Supplementary Table S3.** Cell lines used in this study.

| Name | Genotype | Plasmids used | Resistance marker | Internal Lab ID | Already published or generated for this study |
| --- | --- | --- | --- | --- | --- |
| HeLa Kyoto | Wild type | n/a | n/a | 1 |  |
| HeLa Kyoto SMC4-AID | SMC4-mAID-Halo<br>OsTIR1(F74G)-<br>SNAP-IRES-Blast<br>(integrated into<br>AAVS1 safe<br>harbour locus) | SMC4-mAID-Halo repair template<br>OsTIR1<sup>F74G</sup>-SNAP-IRES-Blast repair template<br>sgRNA expressing/Cas9-human geminin fusion<br>(Cas9-hGem) to tag SMC4<br>sgRNA expressing/Cas9-human geminin fusion<br>(Cas9-hGem) to integrate OsTIR1<sup>F74G</sup> | Blasticidin S<br>Puromycin | 2056 | Published in<br>Schneider et al;<br>2022 |
| HeLa Kyoto NIPBL-AID | mEGFP-mAID-NIPBL<br>OsTIR1(F74G)—<br>SNAP-IRES-Blast<br>(integrated into<br>AAVS1 safe<br>harbour locus) | mEGFP-mAID-NIPBL repair template<br>OsTIR1<sup>F74G</sup>-SNAP-IRES-Blast repair template<br>sgRNA expressing/Cas9-human geminin fusion<br>(Cas9-hGem) to tag NIPBL<br>sgRNA expressing/Cas9-human geminin fusion<br>(Cas9-hGem) to integrate OsTIR1<sup>F74G</sup> | Blasticidin S | 2045 | Generated for<br>this study |
| HeLa Kyoto WAPL-AID | Halo-mAID-WAPL<br>SCC1-mEGFP<br>TIR1-3xMyc-T2A-Puro (Lentiviral<br>integration) | Halo-mAID-WAPL repair template<br>SCC1-mEGFP repair template<br>TIR1-3xMyc-T2A PuroLentivirus<br>sgRNA expressing/Cas9(D10A) to tag WAPL | Puromycin | 1802 | Generated for<br>this study |
| HeLa Kyoto WAPL-FKBP12<sup>F36V</sup> (WAPL-dTAG) | Blasticidin-P2A-2xHA-FKBP12<sup>F36V</sup>(dTAG)<br>-WAPL | Blasticidin-P2A-2xHA-FKBP12<sup>F36V</sup> repair<br>template<br>sgRNA expressing / Cas9<sup>D10A</sup> to tag WAPL | Blasticidin S | 2096 | Generated for<br>this study |
| HeLa Kyoto SMC4-AID_WAPL FKBP12<sup>F36V</sup><br>(WAPL-dTAG) | SMC4-mAID-Halo<br>Blasticidin-P2A-2xHA-FKBP12<sup>F36V</sup>(dTAG)<br>-WAPL<br>OsTIR1<sup>F74G</sup>-SNAP-<br>IRES-Blast<br>(integrated into<br>AAVS1 safe<br>harbour locus) | SMC4-mAID-Halo repair template<br>Blasticidin-P2A-2xHA-FKBP12<sup>F36V</sup> repair<br>template<br>OsTIR1<sup>F74G</sup>-SNAP-IRES-Blast repair template<br>sgRNA expressing/Cas9-human geminin fusion<br>(Cas9-hGem) to tag SMC4<br>sgRNA expressing/Cas9-human geminin fusion<br>(Cas9-hGem) to integrate OsTIR1<sup>F74G</sup> | Blasticidin S<br>Hygromycin<br>B<br>Puromycin | 2108 | Generated for<br>this study, base<br>cell line HeLa<br>Kyoto SMC4-AID cell line<br>mentioned<br>above |

**Supplementary Table S4.** scsHi-C read statistics for ΔNIPBL G2 cells.

| Replicate | Total | Mapped | Unique | Cis | Trans<br>Heterologue | Cis 1Kb+ | Cis 10Kb+ | Cis<br>sister | Trans<br>sister |
| --- | --- | --- | --- | --- | --- | --- | --- | --- | --- |
| Replicate 1 | 6.09E+07 | 5.00E+07 | 4.18E+07 | 3.41E+07 | 7.68E+06 | 2.57E+07 | 1.94E+07 | 2.08E+06 | 4.79E+05 |
| Replicate 2 | 7.93E+07 | 6.30E+07 | 3.53E+07 | 2.85E+07 | 6.85E+06 | 2.13E+07 | 1.58E+07 | 1.51E+06 | 3.49E+05 |
| Replicate 3 | 7.36E+07 | 5.98E+07 | 5.06E+07 | 4.14E+07 | 9.28E+06 | 3.08E+07 | 2.27E+07 | 2.47E+06 | 5.56E+05 |
| Replicate 4 | 8.93E+07 | 7.22E+07 | 5.79E+07 | 4.74E+07 | 1.04E+07 | 3.53E+07 | 2.58E+07 | 2.56E+06 | 5.60E+05 |
| Replicate 5 | 6.55E+07 | 5.37E+07 | 4.28E+07 | 3.49E+07 | 7.92E+06 | 2.61E+07 | 1.93E+07 | 2.00E+06 | 4.59E+05 |
| Replicate 6 | 7.04E+07 | 5.74E+07 | 4.39E+07 | 3.59E+07 | 8.00E+06 | 2.60E+07 | 1.95E+07 | 9.43E+05 | 1.92E+05 |
| Replicate 7 | 6.51E+07 | 5.19E+07 | 4.06E+07 | 3.27E+07 | 7.82E+06 | 2.35E+07 | 1.78E+07 | 6.74E+05 | 1.29E+05 |
| Replicate 8 | 6.14E+07 | 5.01E+07 | 4.49E+07 | 3.60E+07 | 8.89E+06 | 2.62E+07 | 2.00E+07 | 1.28E+06 | 2.77E+05 |
| Replicate 9 | 7.01E+07 | 5.57E+07 | 4.84E+07 | 3.81E+07 | 1.03E+07 | 2.76E+07 | 2.10E+07 | 1.39E+06 | 3.00E+05 |
| Replicate 10 | 5.72E+07 | 4.61E+07 | 4.09E+07 | 3.35E+07 | 7.43E+06 | 2.23E+07 | 1.66E+07 | 7.86E+05 | 1.28E+05 |

**Supplementary Table S5.**
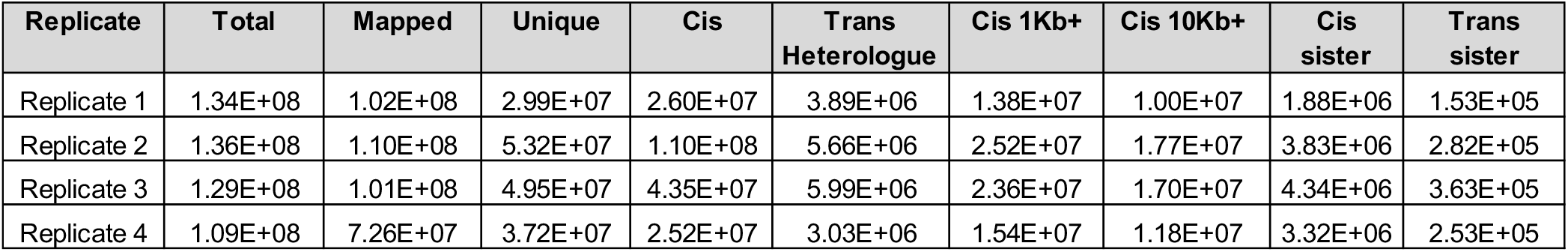
scsHi-C read statistics for ΔSMC4 G2 cells.

| Replicate | Total | Mapped | Unique | Cis | Trans<br>Heterologue | Cis 1Kb+ | Cis 10Kb+ | Cis<br>sister | Trans<br>sister |
| --- | --- | --- | --- | --- | --- | --- | --- | --- | --- |
| Replicate 1 | 1.34E+08 | 1.02E+08 | 2.99E+07 | 2.60E+07 | 3.89E+06 | 1.38E+07 | 1.00E+07 | 1.88E+06 | 1.53E+05 |
| Replicate 2 | 1.36E+08 | 1.10E+08 | 5.32E+07 | 1.10E+08 | 5.66E+06 | 2.52E+07 | 1.77E+07 | 3.83E+06 | 2.82E+05 |
| Replicate 3 | 1.29E+08 | 1.01E+08 | 4.95E+07 | 4.35E+07 | 5.99E+06 | 2.36E+07 | 1.70E+07 | 4.34E+06 | 3.63E+05 |
| Replicate 4 | 1.09E+08 | 7.26E+07 | 3.72E+07 | 2.52E+07 | 3.03E+06 | 1.54E+07 | 1.18E+07 | 3.32E+06 | 2.53E+05 |

**Supplementary Table S6.** scsHi-C read statistics for ΔWAPL G2 cells.

| Replicate | Total | Mapped | Unique | Cis | Trans Heterologue | Cis 1Kb+ | Cis 10Kb+ | Cis sister | Trans sister |
| --- | --- | --- | --- | --- | --- | --- | --- | --- | --- |
| Replicate 1 | 3.50E+07 | 2.68E+07 | 1.64E+07 | 1.42E+07 | 2.25E+06 | 9.84E+06 | 7.64E+06 | 8.09E+05 | 1.06E+05 |
| Replicate 2 | 4.64E+07 | 3.12E+07 | 9.40E+06 | 7.85E+06 | 1.55E+06 | 4.84E+06 | 3.80E+06 | 4.66E+05 | 5.27E+04 |
| Replicate 3 | 5.62E+07 | 3.13E+07 | 1.15E+07 | 9.77E+06 | 1.70E+06 | 6.46E+06 | 5.02E+06 | 5.31E+05 | 6.16E+04 |
| Replicate 4 | 7.99E+07 | 6.17E+07 | 1.02E+07 | 9.09E+06 | 1.11E+06 | 6.07E+06 | 4.54E+06 | 6.95E+05 | 9.36E+04 |
| Replicate 5 | 1.00E+08 | 7.03E+07 | 1.92E+07 | 1.69E+07 | 2.32E+06 | 1.24E+07 | 9.86E+06 | 9.28E+05 | 1.53E+05 |
| Replicate 6 | 9.30E+07 | 7.01E+07 | 8.50E+06 | 7.41E+06 | 1.09E+06 | 4.91E+06 | 3.78E+06 | 4.92E+05 | 6.75E+04 |

**Supplementary Table S7.**
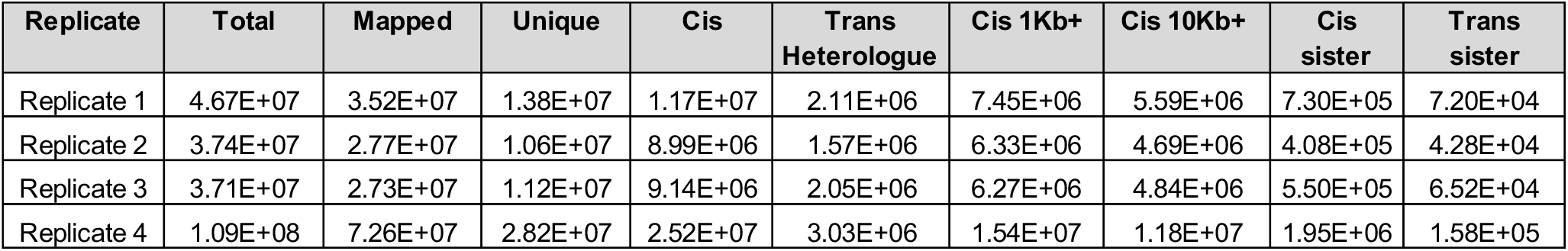
scsHi-C read statistics for ΔWAPL ΔSororin G2 cells.

**Supplementary Table S8.**
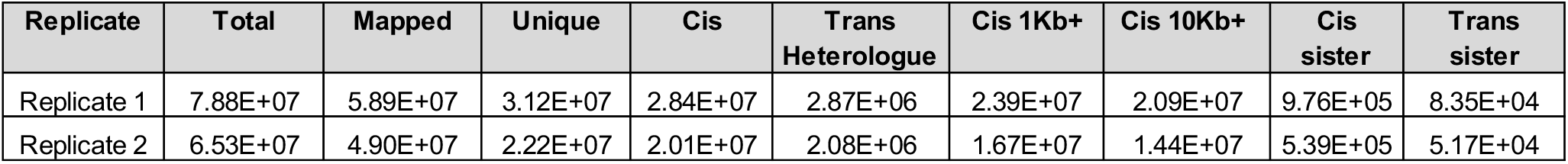
scsHi-C read statistics for wild type prometaphase cells.

| Replicate | Total | Mapped | Unique | Cis | Trans Heterologue | Cis 1Kb+ | Cis 10Kb+ | Cis sister | Trans sister |
| --- | --- | --- | --- | --- | --- | --- | --- | --- | --- |
| Replicate 1 | 7.88E+07 | 5.89E+07 | 3.12E+07 | 2.84E+07 | 2.87E+06 | 2.39E+07 | 2.09E+07 | 9.76E+05 | 8.35E+04 |
| Replicate 2 | 6.53E+07 | 4.90E+07 | 2.22E+07 | 2.01E+07 | 2.08E+06 | 1.67E+07 | 1.44E+07 | 5.39E+05 | 5.17E+04 |

**Supplementary Table S9.** scsHi-C read statistics for ΔNIPBL prometaphase cells.

| Replicate | Total | Mapped | Unique | Cis | Trans Heterologue | Cis 1Kb+ | Cis 10Kb+ | Cis sister | Trans sister |
| --- | --- | --- | --- | --- | --- | --- | --- | --- | --- |
| Replicate 1 | 3.65E+07 | 2.85E+07 | 7.66E+06 | 6.48E+06 | 1.18E+06 | 3.80E+06 | 3.15E+06 | 2.49E+05 | 2.47E+04 |
| Replicate 2 | 4.20E+07 | 3.09E+07 | 1.45E+07 | 1.28E+07 | 1.76E+06 | 9.02E+06 | 7.69E+06 | 5.03E+05 | 4.62E+04 |
| Replicate 3 | 2.78E+07 | 2.08E+07 | 6.71E+06 | 5.61E+06 | 1.10E+06 | 4.33E+06 | 3.74E+06 | 1.74E+05 | 2.11E+04 |

**Supplementary Table S10.** scsHi-C read statistics for ΔSMC4 prometaphase cells.

| Replicate | Total | Mapped | Unique | Cis | Trans Heterologue | Cis 1Kb+ | Cis 10Kb+ | Cis sister | Trans sister |
| --- | --- | --- | --- | --- | --- | --- | --- | --- | --- |
| Replicate 1 | 7.56E+07 | 5.40E+07 | 3.54E+07 | 2.82E+07 | 7.24E+06 | 2.36E+07 | 2.00E+07 | 1.38E+06 | 4.72E+05 |
| Replicate 2 | 7.38E+07 | 5.46E+07 | 4.34E+07 | 3.72E+07 | 6.14E+06 | 2.75E+07 | 2.18E+07 | 1.96E+06 | 3.79E+05 |

**Supplementary Table S11.** Published datasets used in this study.

| Name | GEO ID | Description |
| --- | --- | --- |
| Hi-C: G2 | GSM4613674 | Cooler files for wild type HeLa Kyoto cells, synchronised to G2 (All contacts, cis sister contacts, trans sister contacts) |
| Hi-C: Sororin-AID + auxin G2 | GSM4613677 | Cooler files for Sororin depleted HeLa Kyoto cells, synchronised to G2 (All contacts, cis sister contacts, trans sister contacts) |

